# Distinct plasmid- and host-encoded mechanisms drive small plasmid copy number-mediated heteroresistance in *Escherichia coli*

**DOI:** 10.64898/2026.08.19.745753

**Authors:** Angelika Ntokaki, Enrique Joffré, Bart Melgers, Margarita Komi, Erik Holmqvist, Dan I. Andersson, Hervé Nicoloff, Helen Wang

## Abstract

Antibiotic heteroresistance, the presence of a rare resistant subpopulation within an otherwise susceptible bacterial population, poses a significant clinical challenge. Understanding its genetic mechanisms is critical for early detection and treatment efficacy. Here, we investigate the contribution of small plasmids to heteroresistance using a clinical bloodstream *Escherichia coli* isolate carrying a 12 kb ColE1-type plasmid. We show that this plasmid drives transient β-lactam heteroresistance through massive increases in plasmid copy number. Two distinct genetic mechanisms drive this amplification: mutations in the plasmid RNAI/RNAII that deregulate replication control, and a chromosomal *recD* mutation that induces multimerization and a shift toward rolling-circle replication. Notably, this *recD*-mediated amplification is restricted to small ColE1 and F-plasmids. This study highlights the crucial role of small plasmids in resistance evolution, demonstrating that they can cause this phenotype via alternative genetic pathways.

## INTRODUCTION

Antimicrobial resistance is a critical global health threat, with an increasing number of deaths associated with or directly attributable to it^1^. Standard antimicrobial susceptibility testing (AST) evaluates the bacterial population as a whole and often misses phenotypic heterogeneity caused by persistence and heteroresistance (HR). HR is a phenomenon where a small resistant subpopulation exists within a main susceptible population ^2,3^. Described across diverse bacterial species and antibiotic classes^4–7^ HR complicates minimum inhibitory concentration (MIC) interpretations and frequently leads to prolonged and failed treatment due to expansion of the resistant subpopulations under antibiotic treatment^8–10^.

Multiple HR mechanisms lead to either stable or transient resistance^2^. The most common mechanism in Gram-negative bacteria is tandem gene amplification, driven by homologous recombination between direct repeat sequences (e.g., IS elements) flanking an antibiotic resistance gene with a low activity or expression level and unable to confer resistance at a single copy^11–13^. In *Escherichia coli* and other Enterobacteriaceae, tandem gene amplifications typically occur on large, low-copy plasmids, on which the resistance genes and repeat sequences are present^14,15^. These amplifications transiently increase the gene dosage and, as a result, the MIC, but they impose substantial fitness cost and rapidly revert in the absence of antibiotic selection^16,17^. Another transient HR mechanism involves an increased copy number of resistance plasmids^14^. This increase occurs without detectable mutations that disrupt plasmid replication control, and it imposes substantial fitness costs owing to the burden of carrying additional plasmid DNA, making the phenotype unstable. In addition, transposition of resistance genes onto small, cryptic plasmids, followed by an increase in their copy number, was also described as another transient HR mechanism^14^.

Historically, small, cryptic plasmids have long been overlooked as selfish DNA elements because they lack obvious host-beneficial genes^18,19^. They can replicate via different mechanisms, such as theta (θ) and sigma (σ) rolling-circle replication^20^. However, their widespread distribution in environmental^18^ and human-associated^19^ microbiomes, together with their ability to horizontally transfer antibiotic-resistant genes^21,22^, has renewed interest in their potential contribution to antibiotic resistance evolution^14,23^.

In this study, we investigate p12, a cryptic ColE1-type plasmid isolated from a bloodstream infection *E. coli* that acquired a *bla*_TEM-1_ gene through Tn*3* transposition, thereby conferring HR to piperacillin-tazobactam (TZP). We show that p12 drives HR through massive copy number increases mediated by two distinct mechanisms: mutations in the plasmid RNAI/RNAII replication control region and loss-of-function mutations in the chromosomal *recD* gene. RecD inactivation promotes plasmid multimerization and a shift toward rolling-circle replication^24,25^, specifically affecting small plasmids from diverse replicon types but not large plasmids. These findings identify small plasmids as autonomous drivers of HR and provide the first link between RecD function and PCN-mediated HR.

## RESULTS

### p12 belongs to a widely distributed family of small plasmids

The clinical *E. coli* isolate DA62886 carrying a single small plasmid, p12 (11,744 bp), was isolated from a patient with a bloodstream infection in the Uppsala University Hospital, Sweden^26^. This plasmid is comprised of three major functional regions: an ARG (antibiotic resistance gene) cluster spanning 32.2% of the sequence (*bla*_TEM-1,_ *aph(6)-Id, aph(3’’)-Ib, dfrA14* and *sul2*); mobile genetic elements spanning 34.9%; including a Tn3-family transposon (*tnpA* and *tnpR*) and an IS91-family transposase; and a 20.0% backbone region containing the replication origin, RNAI/RNAII replication control elements and the origin of transfer (Fig. 1A).

**Figure 1.**
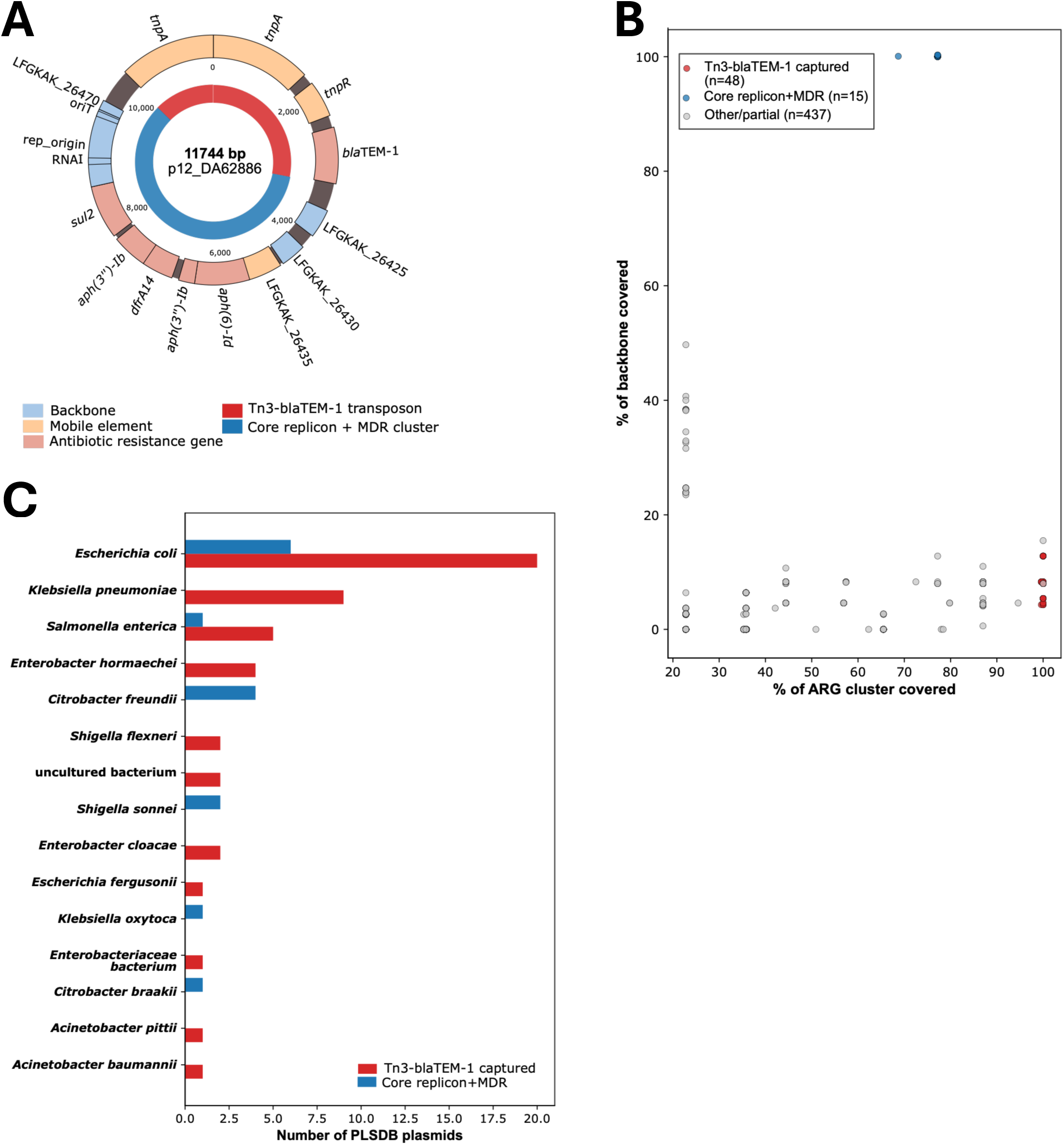
Comparative genomic analysis of plasmid p12. (A). Circular map of p12_DA62886 (11,744 bp). The inner ring indicates the two functionally distinct modules identified by comparison against PLSDB: the Tn3-*bla*_TEM-1_ transposon (red; found in 48 PLSDB plasmids across 11 species) and the core replicon and multidrug-resistance (MDR) cluster (blue; found in 15 PLSDB plasmids across 6 species, all sharing MOB-suite replicon cluster rep_cluster_2335 with p12). The middle ring shows individual annotated features colored by functional category: antimicrobial resistance genes (ARG, red), mobile genetic elements (orange), and backbone/replication-associated features (blue), with gene names indicated at the outer edge. (B). Percentage coverage of the ARG cluster (x-axis) versus the backbone region (y-axis) for each of the 500 PLSDB plasmids sharing homology with p12, derived from BLASTn alignment coordinates. Points are colored by match profile, i.e., hits capturing the Tn3-*bla*_TEM-1_ transposon and full ARG cluster with minimal backbone (red), hits sharing the core replicon and MDR cluster without the *bla*_TEM-1_ insertion (blue), near-full-length matches (green), and other/partial matches (gray). The two-colored point clusters correspond to the two distinct dissemination patterns. (C). Host species distribution of PLSDB plasmids classified as sharing the Tn3-*bla*_TEM-1_ transposon module (red) versus the core replicon and MDR module (blue), illustrating the broader host range of the transposon-borne resistance module relative to the native plasmid replicon.

A BLASTn search against the NCBI non-redundant/nucleotide sequences identified a plasmid (CP164952.1, *E. coli* strain OXEC-283) with 100% identity and coverage across the full 11,744 bp length, isolated from the John Radcliffe Hospital in Oxford, United Kingdom. A BLASTn search against the PLSDB plasmid database (v2024_05_31_v2; 72,556 records), on the other hand, identified 500 plasmids with homology to p12, demonstrating that p12-related sequences are widely distributed. Among the 169 matches covering ≥55% of p12, representing 26 species and 10 genera, comparison of the ARG, mobile-element and backbone regions revealed two distinct dissemination patterns (Fig. 1B). First, the Tn3-*bla*_TEM-1_ transposon together with the full ARG cluster, but little of the p12 backbone (<20% coverage), was found in 48 plasmids across 11 species and 10 replicon incompatibility (Inc) types. These included plasmids from non-Enterobacteriaceae, such as *Acinetobacter* spp., and were considerably larger than p12 (60,296 kb), indicating broad horizontal dissemination of the resistance region.

Second, the native p12 replicon and multidrug-resistant cluster without the *bla*_TEM-1_ insertion (defined by ≥90% backbone coverage, ≥60% ARG coverage and <25% mobile-element coverage) was found in 15 plasmids of similar size (6.7 - 39 kb) across 6 species: *E. coli*, *Salmonella enterica*, *Shigella sonnei*, *Citrobacter freundii*, *Citrobacter braakii, Klebsiella oxytoca*. All belonged to the same MOB-suite replicon cluster as p12 (rep_cluster_2335; Fig. 1C). Together, these findings indicate that p12 represents a prevalent and widely distributed family of small plasmids and that its backbone and resistance regions have followed distinct evolutionary trajectories in different host strains.

### Increased p12 copy number generates TZP HR in *E. coli*

Since p12 carries multiple ARGs, we assessed the susceptibility of the clinical isolate and an MG1655 derivative carrying the plasmid (DA82535) to several antibiotics (Supplementary Dataset 1). HR analysis focused on antibiotics to which the strains were initially susceptible. Both strains exhibited an HR phenotype to piperacillin-tazobactam (TZP; *bla*_TEM-1_), a β-lactam/β-lactamase inhibitor combination frequently used to treat bloodstream infection (Fig. 2A and Fig. 2B), but not to streptomycin (STR; *aph(6)-Id and aph(3”)-Ib*), for which no HR was detected. (Supplementary Fig. 1A and Fig. 1B). TZP-resistant clones were isolated at 2x, 4x, 8x, and 16x the MIC of the main susceptible population, and their p12 copy numbers were quantified by droplet digital PCR (ddPCR). The resistant clones showed a substantial increase in p12 copy number, which naturally exists at ≈ 10 copies per cell, reaching up to a 50-fold increase in the clinical isolate (Fig. 2C) and 200-fold in the MG1655 background (Fig. 2D). These increases elevated the dosage of *bla*_TEM-1_, directly linking p12 amplification to the TZP-HR phenotype. In contrast, p12 plasmid copy number showed little to no variation in the non-STR-HR strains (Supplementary Fig. 1C and Fig. 1D). Thus, a widely distributed small plasmid can generate clinically relevant HR following the acquisition and increased gene copy number (GCN) of *bla*_TEM-1_.

**Figure 2.**
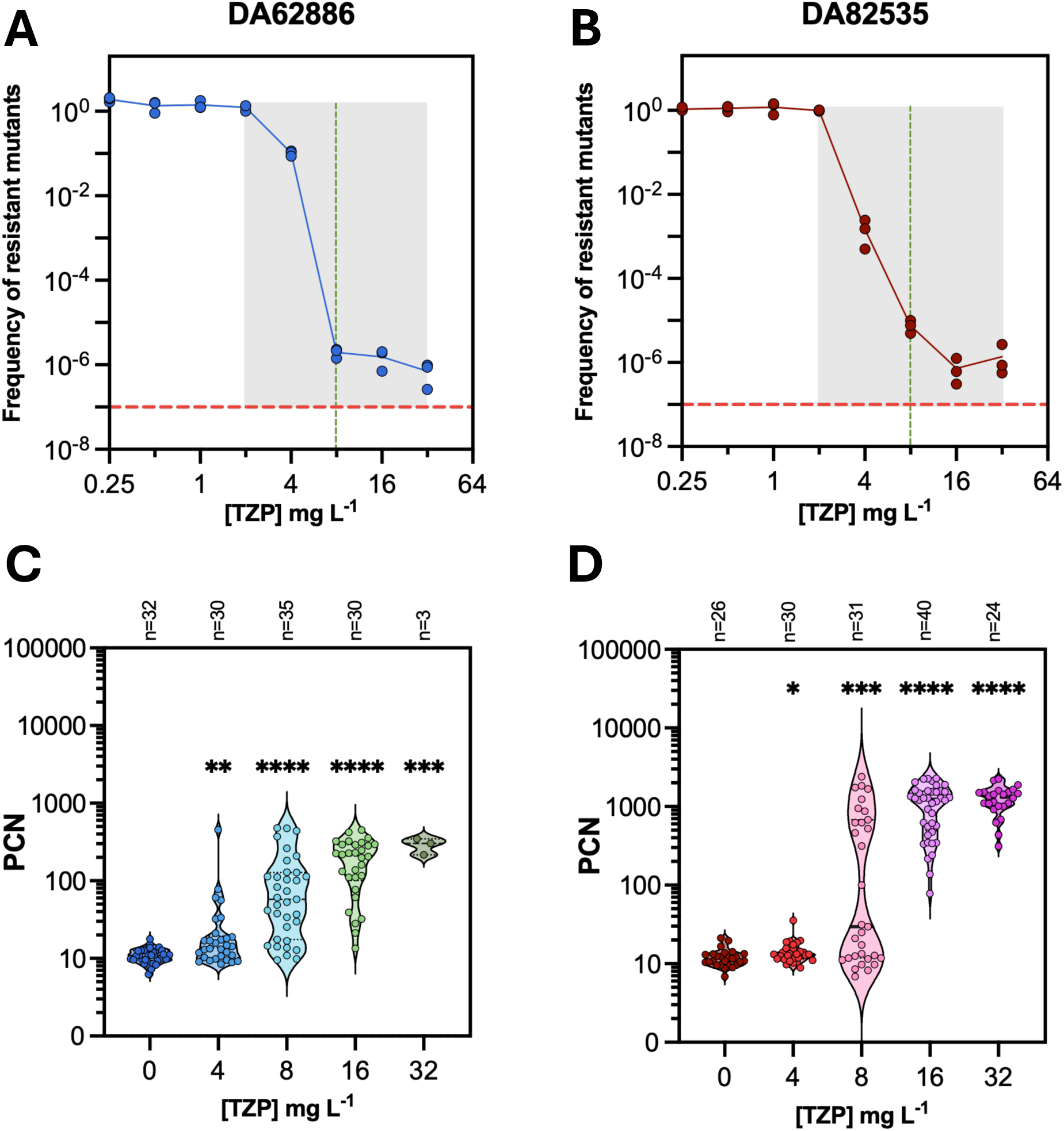
Plasmid p12 carrying one copy of *bla_TEM-1_* generates HR in two *E. coli* genetic backgrounds through PCN increase. (A-B). PAP tests of DA62886 and DA82535, respectively, carrying p12, with gray boxes representing HR phenotypes. Dashed red lines represent the frequency threshold (≥10^-7^), and green lines correspond to the EUCAST clinical breakpoints for TZP (n = 3). (C-D). PCN of isolated resistant mutants at 2x, 4x, 8x, and 16x MIC of the main susceptible population. Data are presented as the median of all replicates. Two-tailed Mann-Whitney tests were performed to compare all groups with the control (0 mg L^-1^ TZP). *: p<0.05, **: p<0.01, ***: p<0.001, ****: p<0.0001

### Two distinct mechanisms drive p12 PCN increase

To identify the mechanisms underlying the observed increased PCN involved in TZP HR, we performed short-read whole-genome sequencing (WGS) of 19 TZP resistant high PCN mutants from each strain background (Table S2). All mutants carried mutations, predominantly in the overlapping RNAI/RNAII replication control region of p12. More specifically, in the clinical isolate DA62886, 74% (14/19) of the mutants carried a 65-bp duplication in the RNAI/RNAII region, whereas the remaining 26% (5/19) carried a single nucleotide polymorphism (SNP) at different positions (Fig. 3A). Among the DA82535 mutants, 63% (12/19) carried duplications; 83% (10/12) had the same 65-bp duplication as the DA62886 mutants and two contained a 38-bp duplication. In all these mutants, the duplicated sequence was detected in only a subset of the population, indicating the coexistence of mutated and wild-type copies of the plasmid in the mutants’ population (Supplementary Fig. 2A). Four mutants carried either a SNP (2/19) or a single nucleotide deletion (InDel) (2/19) in the RNAI/RNAII region. Notably, the remaining three DA82535 mutants (16%) acquired loss-of-function mutations in the chromosomal *recD* gene without any detectable plasmid mutations (Fig. 3B).

**Figure 3.**
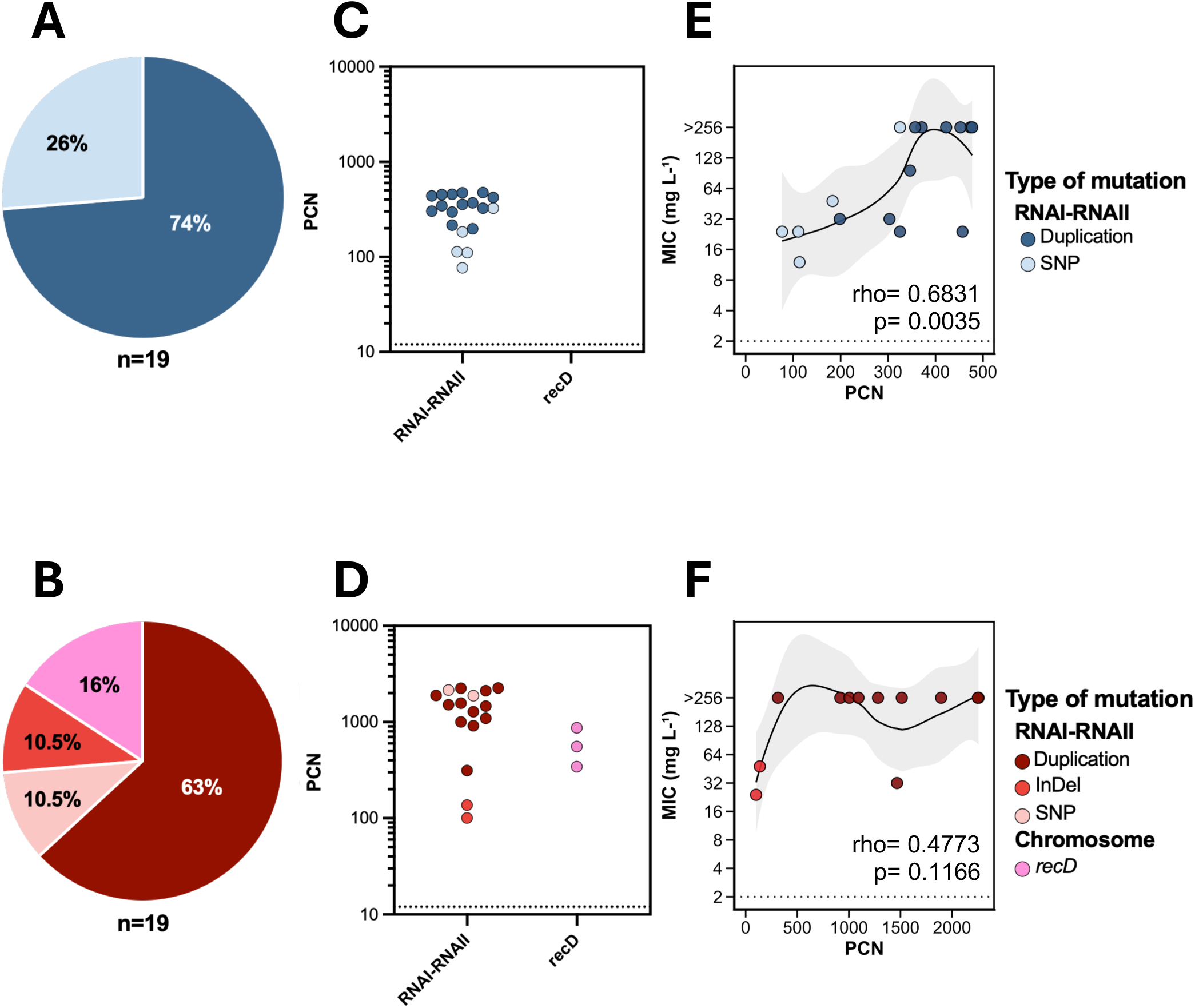
Two mechanisms generate the increase in PCN, leading to higher TZP resistance levels. (A-B). Pie charts showing the prevalence of each mutation type in percentages, per parental strain. **(C-D).** PCN values of resistant mutants of DA62886 and DA82535, respectively, carrying different types of mutations, as represented by the different colors (n = 19). Black dotted line represents the PCN of the respective parental. **(E-F).** Minimum inhibitory concentrations of resistant mutants carrying only one mutation type (n = 16 and n = 12, respectively), in relation to their PCN values. The black line represents the best-fit line of all the data points, the grey area the 95 % CI, and the black dotted line represents the MIC value of the parental strain. A Spearman correlation test was performed in RStudio, and the rho and p - values are shown in the bottom right corner of each graph.

In the clinical DA62886 background, RNAI/RNAII duplications resulted in higher PCN increases than SNPs (Fig. 3C), whereas duplications and SNPs generated similar PCN levels in DA82535. The smallest increase was observed in DA82535 mutants carrying the InDel, which maintained around 100 copies of p12 per cell (Fig. 3D). Collectively, these findings identify two mutually exclusive routes to p12 amplification: disruption of plasmid-encoded RNAI/RNAII replication control and loss of host RecD function.

### Increased p12 copy number is unstable without antibiotic selection

To assess the phenotypic and genotypic stability of the p12 PCN increase, representative mutants from each mutation class were grown for 40 generations without selective pressure. Single revertant clones were then analyzed for p12 PCN by ddPCR and for TZP MIC by E-tests. Representative revertant clones then underwent WGS. PCN values generally decreased in the absence of antibiotics, although they rarely returned to the levels of the original HR strain within 40 generations (Supplementary Fig. 3A and Fig. 3B). In DA62886, mutants carrying SNPs exhibited a significantly slower reversion rate compared to those carrying the 65-bp duplication, consistent with their corresponding changes in TZP resistance levels (Supplementary Fig. 3C). In DA82535, mutants carrying InDel reverted significantly more slowly than the other mutation classes, and many showed no detectable reduction in PCN. Mutants carrying duplications or SNPs displayed a faster reversion rate, whereas *recD* mutants showed the greatest variability. Some *recD* revertants retained the elevated PCN, while 25% of the 54 clones tested (among the original three resistant mutants) completely lost the p12 plasmid (Table S3). Correspondingly, their TZP MICs decreased markedly, in some cases to below the parental MIC, further supporting the p12 plasmid loss (Supplementary Fig. 3D).

WGS of selected revertants (Table S4) showed that reversion was generally not caused by restoration of the original mutation to the wild-type sequence. Instead, among mutants carrying the RNAI/RNAII duplications, the proportion of plasmid molecules containing the duplication decreased during passaging. Some revertants shifted from a duplication-dominated to SNP-dominated population, suggesting that the original mutants might contain a subpopulation of p12 SNP variants at a frequency below the detection limit. As the cultures were grown without selection, these competing plasmid variants changed in relative abundance. A small number of resistant mutants (n = 3 and n = 7 for the two backgrounds, respectively) carried additional mutations, mainly in genes of unknown function. Because these mutations could independently affect the MIC or fitness, these mutants were excluded from further analyses.

### Increased p12 copy number is not consistently associated with proportional increases in MIC or fitness cost

To determine whether increased p12 copy number translated into higher resistance, we measured TZP resistance levels by E-tests for the mutants carrying a single type of mutation. In the DA62886 background, PCN correlated positively with MIC values (Spearman’s rho = 0.6831, p = 0.0035; Fig. 3E), consistent with increased *bla*_TEM-1_ gene dosage contributing to higher MIC. However, this relationship plateaued at higher copy numbers. In mutants derived from DA82535, the association was positive but not significant (Spearman’s rho = 0.4773, p = 0.1166; Fig. 3F), with most mutants reaching maximum measured MIC. Thus, an increase in p12 PCN increased TZP resistance within a limited range, beyond which further increases in PCN produced little detectable change in MIC.

Increased PCN results in a substantial increase in total DNA content per bacterial cell. DA62886-derived mutants carried an estimated 1 to 5.5 Mbp of additional plasmid DNA, in some cases approximately doubling the DNA content relative to the 5.31 Mbp parental genome. Despite this burden, none showed a significant growth defect. Eight of 16 mutants exhibited growth-rate increases of up to 3%, whereas the remainder showed reductions of no more than 3% (Supplementary Fig. 4A).

The DNA burden was even greater in the MG1655 background. DA82535-derived mutants, with PCN increases of up to 200-fold, carried an estimated 1 to 25 Mbp of additional plasmid DNA, corresponding in some cases to approximately five times the parental genome. Ten of 15 mutants (67%) displayed fitness costs of ≥6%, reaching a maximum of 14% (Supplementary Fig. 4B). After 40 generations without selection, growth rates tended to recover in mutants that initially exhibited a fitness cost, whereas they remained stable in the other lineages (Supplementary Fig. 5).

To evaluate the effect of carrying additional DNA, we normalized the change in relative fitness to the amount of additional DNA, where negative values indicate a cost associated with it and positive values indicate a fitness benefit. As illustrated in Supplementary Fig. 4D, mutants carrying an InDel, which had the smallest PCN increase, displayed the highest mean fitness benefit per additional kb (+3.4×10^-6^). Duplication mutants imposed the highest costs, reaching up to -7.38×10^-6^ per kb extra DNA. Unexpectedly, in one case (Supplementary Fig. 4C), mutants carrying a SNP were associated with a higher fitness benefit per additional kilobase (kb) (+2.53×10^-6^ to +2.52×10^-5^) compared to those carrying the duplications (-8.55×10^-6^ to +3.72×10^-6^). These findings indicate that the general fitness reduction due to p12 PCN increase depends on both the bacterial strains and the underlying mutation and is not determined solely by the amount of additional DNA.

### RNAI and RNAII exhibit a change in secondary structure and expression levels

Because most plasmid mutations mapped to the overlapping RNAI/RNAII replication control, we examined their predicted effects on the secondary structures of the molecules. As displayed in Supplementary Figure 6, most single-nucleotide mutations are located in the second hairpin loop of RNAI, suggesting that they could affect RNAI/RNAII recognition and/or binding. By contrast, duplications were predicted either to elongate the second hairpin loop or generate a markedly altered RNAI overall structure (Supplementary Fig. 6G and Fig. 6H).

Northern blot analysis was used to assess RNAI and RNAII transcript profiles in the DA62886 clinical background (Fig. 4). We analyzed a mutant carrying the 65-bp duplication (DA84124) and one of its revertant clones, together with a mutant/revertant pair carrying a SNP at the 5’ end of RNAI (DA84140). The parental strain exhibited low RNAI abundance at steady-state PCN. DA84140 displayed a stronger signal at around 100 nucleotides, potentially reflecting increased gene dosage caused by elevated PCN, together with fainter, smaller bands consistent with RNAI processing products (PT-1 and PT-2). In contrast, DA84124 and its revertant displayed only a single, intense band substantially smaller than the native RNAI. This band, estimated to be around 50 nucleotides in length, may represent a stable RNAI processing or degradation product (PT-3). For RNAII, the abundance was low in the parental strain, with faint bands migrating between 400-500 nucleotides. Mutants and revertants exhibited increased abundance of multiple RNAII transcripts ranging from approximately 190 to >500 nucleotides. The two largest detected bands migrated within the size range expected for the reported 555-nucleotide RNAII pre-primer transcript, although their precise sizes could not be resolved using the current experimental setup. The increased RNAII signal may reflect elevated gene dosage resulting from p12 PCN increase and may also reflect altered RNA processing or stability. Together, these findings support a model in which mutations within the RNAI/RNAII region alter RNA structure and transcript profiles, thereby disrupting replication control and causing an increase in p12 copy number.

**Figure 4.**
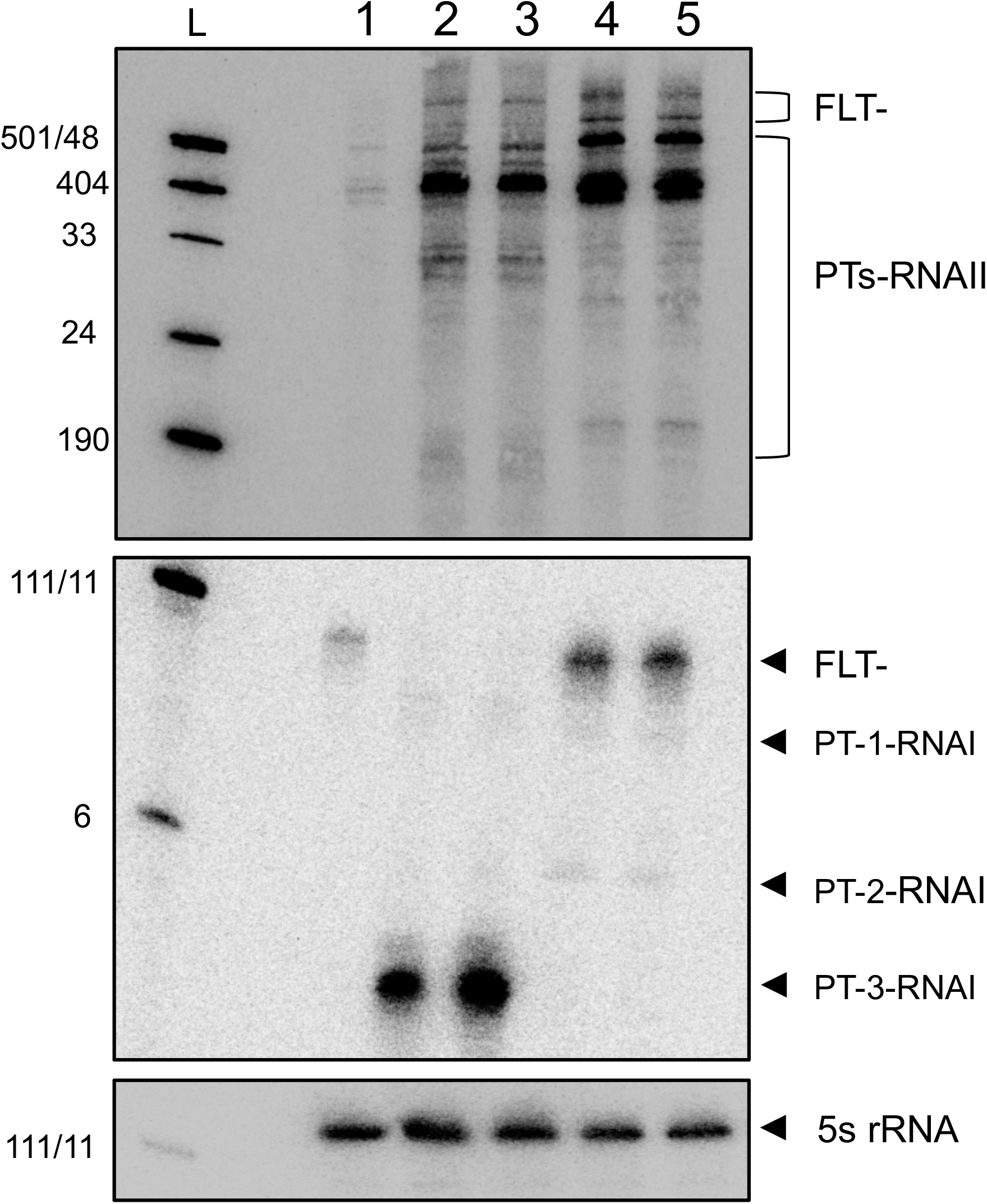
Mutations in the RNAI/RNAII region result in distinct RNAI/RNAII profiles in resistant mutants. Northern blot of DNase-treated RNA samples on a UREA/PAGE gel, labelled with [γ-32P] ATP 5′-end-labeled DNA probes. Lane 1: Parental DA62886, Lane 2: DA84124 (65 bp duplication), Lane 3: DA84124 revertant, Lane 4: DA84140 (5’-SNP), Lane 5: DA84140 revertant. L: pUC19 MSPI marker with the corresponding size of each band. FLT: full-length transcript, PT: processed transcript

### Loss of RecD protein increases p12 copy number and plasmid multimerization

To determine whether the increased p12 copy number associated with *recD* mutations involved plasmid multimerization, we performed Nanopore sequencing of TZP-resistant mutants and mapped the reads to the chromosome and an artificial p12 dimer. Most p12 mapping reads corresponded to the 11,744-bp monomeric plasmid. However, reads exceeding 12 kb and spanning consecutive p12 units were detected in the *recD* mutants DA84222 and DA84157, supporting the presence of plasmid multimers (Supplementary Fig. 7). These mutants and a DA84157 revertant strain exhibited pronounced long right-skewed distributions and the highest proportions of multimeric reads: 4% and 6% in DA84222 and DA84157, respectively, and 5% in the revertant, compared to <0.25% in the parental strain and all other mutants carrying plasmid replication control mutations (Table S2 and Supplementary Figure 8).

To investigate whether this phenotype was specific to RecD or reflected broader disruption of the RecBCD recombination pathway, we utilized Δ*recA,* Δ*recB,* Δ*recC,* and Δ*recD* mutants (without the kanamycin cassette) from the *E. coli* KEIO collection^27^, together with the corresponding wild-type strain. Each strain was transformed with the p12 plasmid. The Δ*recA,* Δ*recB* and Δ*recC* strains showed intrinsic growth defects, regardless of p12 carriage, whereas the Δ*recD* strain grew similarly to the wild-type strain (Supplementary Fig. 9). Moreover, MIC testing of the strains through E-test revealed a different phenotype in the Δ*recD* background carrying p12, with colonies growing within the inhibition zone (Supplementary Fig. 10).

Population analysis profiling revealed TZP HR in all p12-containing strains. However, the Δ*recD* strain displayed a rightward shift of the population profile, indicating an increased resistance of the main population (Fig. 5A). Colonies isolated without antibiotic and at 8x the parental MIC_TZP_ (8xMIC) were quantified for p12 PCN. In the absence of TZP, Δ*recB* and Δ*recC* mutants exhibited approximately a 2-fold increase in PCN compared to the wild-type. Following TZP selection, PCN increased by approximately 20- to 40-fold in all backgrounds except Δ*recD*. Notably, p12 copy number in the Δ*recD* was already nearly 45-fold higher than in the wild-type strain, even in the absence of selection, and increased only a further 1.5-fold at 8xMIC (Fig. 5B). Thus, loss of RecD function was sufficient to generate a constitutively high p12 copy number in the absence of antibiotic exposure, thereby increasing *bla*_TEM-1_ gene dosage and explaining the elevated TZP resistance.

**Figure 5.**
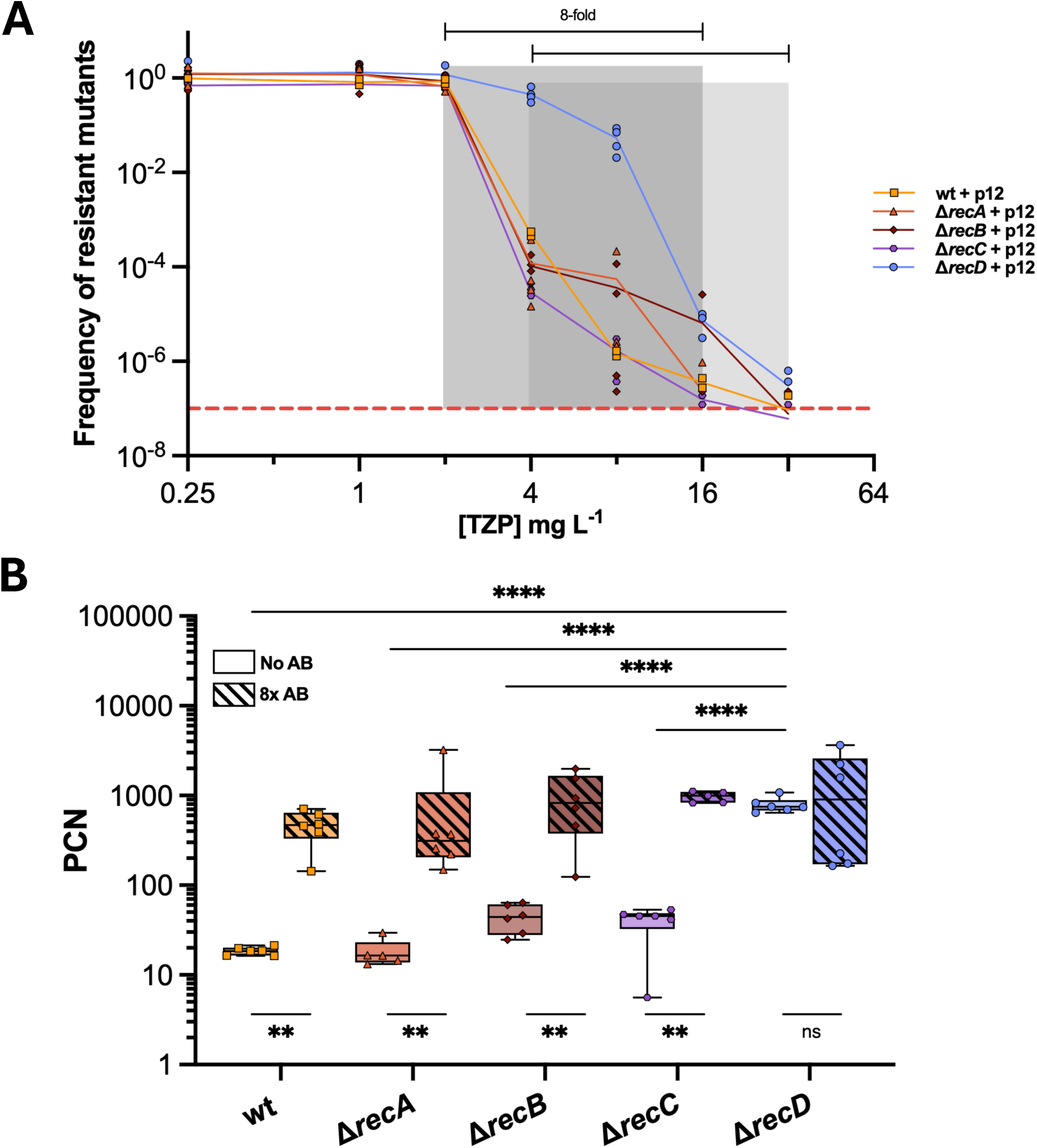
Mutation in the *recD* gene generates HR through increased p12 PCN. (A). TZP PAP tests of KEIO collection mutants Δ*recA*, Δ*recB*, Δ*recC*, Δ*recD*, plus wild-type, carrying p12 (n = 3). Gray boxes represent the HR phenotype, with Δ*recD +* p12 exhibiting a shift in the HADNAG and the overall curve. The dashed red line represents the frequency threshold (≥10^-7^). (B). PCN values of clones isolated from plates with no antibiotic and 8x the MIC of the main susceptible population for each strain (8xAB) (n = 6). An ordinary one-way ANOVA test (Tukey’s correction for multiple comparisons) was performed to compare the groups in the “No AB” condition, and multiple Mann-Whitney tests to compare “No AB” and ”8x MIC” within each group *: p<0.05, **: p<0.01, ***: p<0.001, ****: p<0.0001.

Similar to what was observed with spontaneous *recD* mutants, Nanopore sequencing confirmed increased p12 multimerization in the Δ*recD* mutant, with approximately 3% of p12 reads classified as multimers without selection, and 6% after TZP exposure (Table 1 and Figure 6). The proportion of multimeric reads was significantly higher in Δ*recD* than in the wild-type strain under both conditions (no AB: *p* = 0.023; 8xMIC: *p* = 0.049). Additionally, two-way ANOVA revealed a significant effect of genotype (*p* = 3.2×10^-4^) and a genotype-antibiotic interaction approaching significance (*p* = 0.054), suggesting that TZP exposure may further promote multimerization in Δ*recD*. In contrast, Δ*recA,* Δ*recB,* Δ*recC* resembled the wild-type strain, with ≤1% of reads corresponding to multimers (Table 1 and Supplementary Figures 11, 12). Consistent with these findings, undigested plasmid DNA produced a slower-migrating band exclusively in Δ*recD* (Supplementary Fig. 13A), whereas digestion with the single-cutter enzyme *Hind*III resolved the multimer into a single band corresponding to linearized p12 (Supplementary Fig. 13B). Overall, these results support that RecD loss increases steady-state p12 PCN and promotes the formation of long head-to-tail plasmid multimers (Fig. 6D).

**Table 1.** Mutations in the *recD* gene result in increased p12 PCN and plasmid multimerization. Analysis of nanopore long-read sequences for p12, with and without selective pressure. The last column represents the percentage of multimer reads divided by (monomer + multimer) reads only.

| Genetic background | Plasmid | Selection | % of multimers (n = 3) |
| --- | --- | --- | --- |
| KEIO wild-type | p12 | No AB | 0.00 |
|  |  | 8x MIC | 0.01 ± 0.01 |
| $\Delta recA$ | p12 | No AB | 0.00 |
|  |  | 8x MIC | 0.02 ± 0.01 |
| $\Delta recB$ | p12 | No AB | 0.26 ± 0.30 |
|  |  | 8x MIC | 1.04 ± 0.50 |
| $\Delta recC$ | p12 | No AB | 0.14 ± 0.07 |
|  |  | 8x MIC | 0.54 ± 0.13 |
| $\Delta recD$ | p12 | No AB | 2.77 ± 0.74 |
|  |  | 8x MIC | 6.10 ± 2.44 |

**Figure 6.**
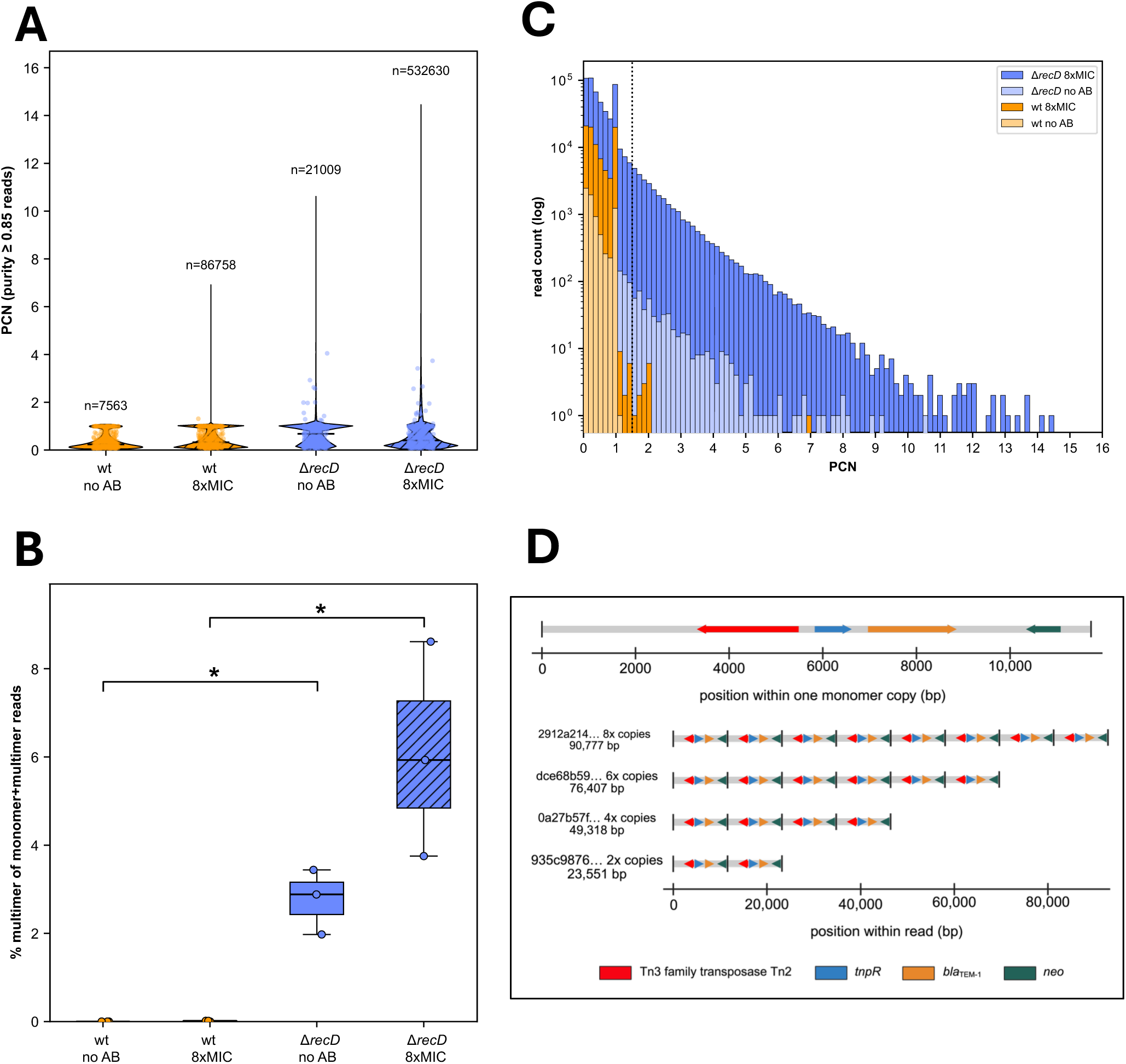
Plasmid copy number and multimerization in KEIO wt versus Δ*recD*, with and without antibiotic selection. Long-read sequencing of plasmid p12 was performed in ΚΕΙΟ wild-type (wt) and Δ*recD E. coli* backgrounds under no antibiotic (no_AB) or 8x MIC (8xAB) selection, with n = 3 biological replicates per genotype/condition. Per-read plasmid copy number (PCN) was estimated as the merged span of all plasmid-mapping alignment intervals divided by the monomer reference length; reads were included in copy number analyses only if ≥85% of their length mapped to the plasmid. Reads with PCN ≥ 1.5 were classified as multimeric, otherwise as monomeric/fragment. (A). Violin plots of the distribution of per-read copy number estimation among purity-filtered reads, pooled across the 3 replicates per group. Individual reads are displayed as semi-transparent points. n above each violin gives the number of reads underlying that distribution. (B). Percentage of plasmid-classified reads scored as multimeric [100 x n(multimer) / (n(monomer) + n(multimer))], calculated per biological replicate (n = 3 per genotype per condition, the true statistical unit) and shown as boxplots (median, interquartile range, min–max whiskers) with individual replicate values overlaid. Comparison between Δ*recD* and wt within each antibiotic condition was done using Welch’s t-test and two-sided Mann-Whitney U test, and two-way ANOVA was used to compare genotype vs antibiotic exposure. *: p<0.05 (C). Overlaid histograms of per-read copy number estimation (purity-filtered reads, pooled across replicates) for all four genotype-per-condition groups. The dotted line marks the PCN = 1.5 multimer threshold. (D). Gene map of the p12 plasmid generated from the Bakta annotation. Protein-coding sequences are colored according to gene identity (top panel). Representative Oxford Nanopore reads from the *recD* mutant classified as plasmid multimers, containing two to eight tandem head-to-tail copies of the p12 plasmid, demonstrating that individual long reads span complete tandem repeats of the plasmid.

### RecD-dependent PCN increase is specific to small plasmids

Since RecD-dependent plasmid multimerization has previously been described for ColE1 plasmids^25^, we tested whether the effect extended to plasmids of various sizes and replication types. In addition to p12, we examined p3, a small ColE1-type plasmid isolated from a clinical strain carrying the fluoroquinolone resistance gene *qnrB19*; a 9-kb miniF (IncFIA) plasmid; and two large IncF plasmids, p139 (CP029577) and p178 (CP029580). PlasmidFinder 2.0 identified that the large plasmids contain more than one origin of replication (Supplementary Fig. 14).

Relative to the wild-type background, deletion of *recD* increased p3 copy number significantly, by approximately 35- to 50-fold (Fig. 7A), and miniF copy number by approximately 30- to 100-fold (Fig. 7B). In contrast, the PCN of the large IncF plasmids was unaffected (Fig. 7C and Fig. 7D). Thus, RecD-dependent PCN increase is not restricted to ColE1 replicons but is specific to small plasmids, whereas the copy number of large plasmids remains unchanged. Moreover, we examined the RecD-dependent multimerization in strains carrying p3- and miniF (Table 2). Nanopore sequencing analysis revealed negligible multimerization for both plasmids in the wild-type, Δ*recA,* Δ*recB,* and Δ*recC* (<0.5%), whereas multimeric reads increased to 6% for miniF and 39% for p3 in Δ*recD* (Table 2 and Supplementary Figures 15, 16). These results highlight that RecD loss promotes multimerization across small plasmids with distinct replication systems, including both ColE1- and F-like replicons.

**Figure 7.**
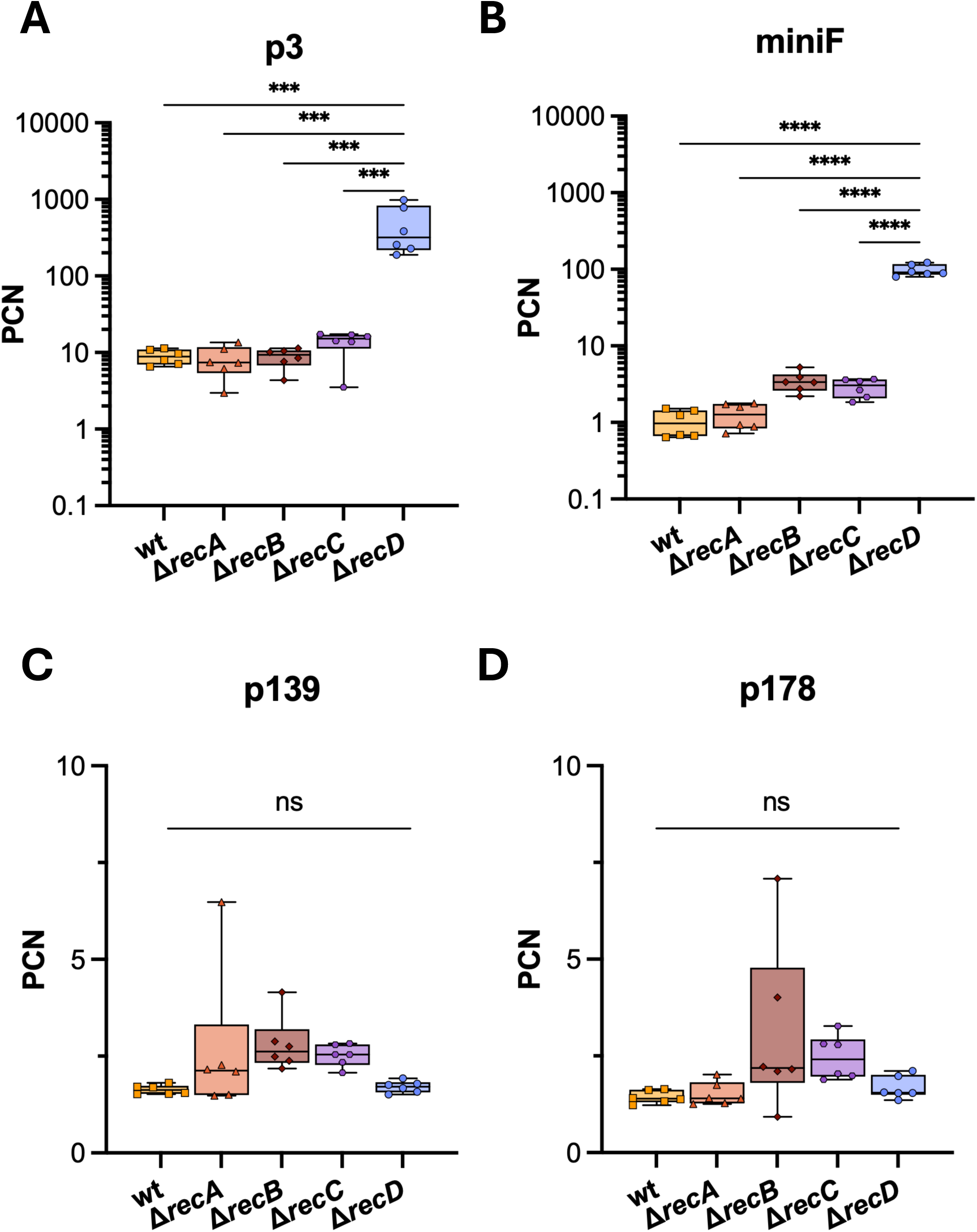
Mutation in the *recD* gene results in increased PCN for more small plasmids. (A-D). Steady-state PCN of four plasmids - p3, miniF, p139, and p178, respectively – in the five KEIO genetic backgrounds (n = 6), analyzed with ddPCR. One-Way ANOVA tests with multiple comparisons and Tukey’s post hoc correction were performed in GraphPad Prism 11, after having tested for normality of the data with a Shapiro-Wilk test. ns: non-significant, *: p<0.05, **: p<0.01, ***: p<0.001, ****: p<0.0001.

**Table 2.** Mutations in the *recD* gene result in increased PCN and plasmid multimerization for more small plasmids. Analysis of nanopore long-read sequences for p3 (plus more small plasmids carried in Δ*recC* and Δ*recD* backgrounds – p2 and p5), and miniF, without selective pressure. The last column represents the percentage of multimer reads divided by (monomer + multimer) reads only.

| Genetic background | Plasmid | Selection | % of multimers (n = 3) |
| --- | --- | --- | --- |
| KEIO wild-type | p3 | No AB | 0.16 ± 0.19 |
| $\Delta recA$ | p3 | No AB | 0.41 ± 0.16 |
| $\Delta recB$ | p3 | No AB | 0.23 ± 0.06 |
| $\Delta recC$ | p3 | No AB | 0.05 ± 0.04 |
| $\Delta recD$ | p3 | No AB | 38.65 ± 1.01 |
| KEIO wild-type | pminiF | No AB | 0.00 |
| $\Delta recA$ | pminiF | No AB | 0.03 ± 0.04 |
| $\Delta recB$ | pminiF | No AB | 0.58 ± 0.35 |
| $\Delta recC$ | pminiF | No AB | 0.20 ± 0.35 |
| $\Delta recD$ | pminiF | No AB | 5.97 ± 2.22 |

## DISCUSSION

Accurate detection of HR and an understanding of its mechanisms are critical for guiding antimicrobial treatment^26^. In this study, we examined the role of previously neglected small plasmids as drivers of HR evolution. We show that small clinical plasmids can autonomously promote HR by increasing plasmid copy number and subsequently resistant-gene dosage. Using the small plasmid p12 in both its native bloodstream infection *E. coli* isolate and an MG1655 background, we identified two distinct routes to PCN increase: disruption of the plasmid-encoded RNAI/RNAII replication control system and loss of chromosomal RecD function. These findings expand the known mechanisms of gene-dosage-mediated HR and identify small plasmids as active drivers of transient antibiotic heteroresistance.

Increase in p12 PCN generated HR toward TZP in both genetic backgrounds, although the magnitude of the PCN increase differed. PCN increased by up to 50-fold in mutants derived from the clinical isolate, but as much as 200-fold in the MG1655 background, indicating that PCN is influenced by the host genotype. One possible explanation is that the native clinical host possesses regulatory or physiological constraints that limit excessive p12 replication, whereas the naïve MG1655 background lacks these. Moreover, PCN and *bla*_TEM-1_ gene dosage were positively correlated with the TZP resistance levels at lower copy numbers, but this relationship plateaued at higher copy numbers. Thus, increasing β-lactamase dosage results in increased resistance levels up to a certain point, after which increased gene copy number does not lead to further increases in MIC, consistent with previous observations^14^. This plateau may reflect saturation of β-lactamase activity or the export pathway, physiological constraints imposed by extreme gene expression, or limitations in MIC resolution.

Most resistant mutants carried mutations in the overlapping RNAI and RNAII region responsible for ColE1-type plasmid replication control. RNAII acts as the replication primer: it hybridizes with the template DNA and is cleaved by RNase H to a mature ∼550-nucleotide transcript to initiate replication^28^. RNAI is an approximately 110-nucleotide antisense RNA that binds nascent RNAII and prevents formation of the RNA-DNA hybrid required for replication^29,30^. Their initial “kissing-complex” interaction is mediated by complementary stem-loops and subsequently stabilized by Rop^31,32^. The 5′ single-stranded regions are particularly important for initiating this interaction, and even small sequence changes can markedly reduce RNAI–RNAII binding^29,33^. Because RNAI is substantially more abundant than RNAII, modest changes in its levels can have major effects on the activity of RNAII and PCN^34^.

Our Northern blot analysis indicates that mutations in this regulatory region alter the RNA processing or structure rather than acting solely through changes in transcript abundance. The 65-bp duplication is predicted to generate an elongated double-stranded RNA stem-loop that could serve as a substrate for RNase III^35^, consistent with the appearance of a smaller, stable RNAI fragment. The 5′-SNP mutant showed increased full-length RNAI together with low-abundance degradation products. The stronger RNAI signal observed is partly a result of increased gene dosage with higher PCN. Greater RNAI abundance does not necessarily indicate greater inhibitory activity, rather, the RNAI/RNAII ratio dictates the replication rate. Because the SNP lies within the 5′ region involved in pairing initiation, it could alter RNA folding or the kinetics of RNAI-RNAII complex formation, thereby weakening replication control. Indeed, single-nucleotide changes in the RNAI/RNAII hairpins can disrupt RNAI-mediated inhibition by altering RNA structure and intermolecular pairing^36^.

High PCN mutants also accumulated multiple RNAII species, including transcripts longer than 500 nucleotides that were consistent in size with the reported 555-nucleotide RNAII pre-primer. Their increased abundance could result from elevated plasmid dosage. Alternatively, accumulation of longer RNAII transcripts might contribute directly to increased replication, because single-nucleotide changes can influence RNA-DNA hybrid formation, secondary structure and RNase H processing^37^. Although the present data cannot distinguish cause from consequence, they indicate that RNAI/RNAII mutations probably deregulate replication through altered RNA structure, processing, stability or duplex formation rather than through transcript abundance alone. High-resolution mapping of transcript ends, for example by 5′ RACE, will be required to define the underlying molecular changes.

The second amplification route involved loss-of-function mutations in chromosomal *recD*. RecD is a component of the RecBCD complex, which participates in homologous recombination and the processing of replication intermediates^38–41^. Loss of RecD converts RecBCD into a highly recombinogenic helicase with reduced nuclease activity, resembling the state induced when RecBCD encounters a Chi site^42^. Previous studies showed that *recD* mutations promote increases in plasmid copy number and formation of long plasmid multimers in both ColE1 and mini-F systems^25,43,44^. Consistent with these findings, p12 multimerization occurred at low frequency in parental strains but increased substantially following *recD* deletion. Agarose gel electrophoresis also revealed an additional p12 species specifically in the Δ*recD* background, probably representing a multimeric covalently closed or open circular form^43^.

The Δ*recD* effect was specific to particular plasmid classes. Loss of RecD increased the copy number of the clinical ColE1-type plasmids p12 and p3 and of mini-F plasmids, but not that of large IncF plasmids. The phenotype therefore appears to depend on plasmid size, replication mechanism, or both. In addition to its established role in homologous recombination, RecBCD is also required for the proper completion of chromosome replication^45,46^. Our results further showed that *recD* mutations were associated with head-to-tail multimer formation and DNA amplification, suggesting that loss of RecD may permit rolling-circle-like σ replication. In the model proposed by Goswami and Gowrishankar, RecBCD nuclease activity suppresses σ-type over-replication by processing DNA structures generated at converging replication forks^47^. In the absence of RecD, these recombination and replication intermediates may persist, allowing aberrant replication and multimer formation. Although further experiments are required to demonstrate this mechanism directly, our findings provide, to our knowledge, the first link between RecD and PCN-mediated HR.

Increased p12 copy number was generally unstable in the absence of TZP, consistent with the transient nature of amplification-mediated HR^14,48,49^. The rate of PCN decline depended on the underlying mutation: mutants carrying the 65-bp duplication reverted more rapidly than those carrying SNPs, small InDels, or *recD* mutations. Because the duplication was present in only a fraction of plasmid molecules, cells dominated by wild-type p12 variants could rapidly outcompete high-copy variants after removal of antibiotic selection. WGS showed that reversion generally did not result from restoration of the original mutation. Instead, the fraction of p12 molecules carrying the duplication declined during passaging, and some populations shifted from being dominated by the duplication to being dominated by an SNP. These observations indicate that individual resistant cultures contained heterogeneous plasmid populations and that variants with lower costs or more stable replication control were selectively enriched after antibiotic withdrawal.

In MG1655, the 65-bp duplication produced p12 DNA levels equivalent to as many as five chromosome copies yet imposed an average fitness cost of only approximately 15%; this cost was largely eliminated in revertants. The modest cost associated with the extraordinary DNA burden suggests that plasmid fitness costs are only weakly associated with the amount of additional DNA^14^. As shown previously, many plasmid-encoded genes impose little measurable cost^50^, which potentially could arise from specific plasmid-host interactions, including altered activation of chromosomal functions^51^. Interestingly, our findings show that in some cases, p12 amplification can even confer a fitness benefit for the host. Given the high prevalence, widespread distribution, and low fitness costs associated with their amplification, these overlooked small plasmids may serve as important vehicles for the dissemination of HR.

Multimerization may further destabilize p12 inheritance. Small, high-copy plasmids generally lack active partitioning systems and instead rely on their high abundance to ensure random distribution of copies to both daughter cells^52^. Multimerization reduces the number of independently segregating plasmid molecules without necessarily reducing the total amount of plasmid DNA, thereby increasing the probability of generating plasmid-free daughter cells^53^. The complete loss of p12 in a subset of Δ*recD* revertants is consistent with this model and may explain the pronounced reduction in TZP MIC observed in these clones.

The instability of small plasmid p12 PCN has potential diagnostic implications. Resistant subpopulations may rapidly decrease in frequency during growth (e.g., in blood cultures) and routine antimicrobial susceptibility testing, causing HR isolates to be misclassified as susceptible^54^. Small plasmids could therefore contribute to otherwise unexplained treatment failure, particularly because their limited gene content often causes them to be overlooked in genomic and epidemiological analyses. Further clinical studies are needed to determine how frequently this mechanism affects susceptibility testing and treatment outcomes.

Finally, comparative sequence analysis showed that p12 belongs to a conserved, mobilizable small-plasmid lineage, rep_cluster_2335, that has acquired a Tn3-*bla*_TEM-1_ insertion. Whereas the native plasmid backbone circulates predominantly among closely related Enterobacteriaceae, its transposon-borne resistance region has disseminated across a broader range of species and unrelated plasmid backbones. Small plasmids can therefore capture, maintain, and redistribute resistance genes while also enabling their massive transient amplification. This allows them to generate HR autonomously, bypassing the need for large multidrug-resistance plasmids commonly associated with resistance-gene amplification. Collectively, our findings consolidate small plasmids as drivers of HR and suggest that their contribution to the emergence, evolution, and dissemination of antimicrobial resistance is greater than previously suggested.

## MATERIALS AND METHODS

### Bacterial strains, plasmid transformation, media, and growth conditions

All the bacterial strains used in this study are described in Table S1 of the supplementary material. Growth was routinely performed in Mueller-Hilton (Difco, Becton Dickinson Company) broth (MHB) and MH agar (MHA). All incubations were carried out at 37 °C, with agitation (180 rpm) for liquid cultures. The antibiotic stocks were purchased from Sigma-Aldrich and were always prepared fresh, as were the plates used in PAP tests. For a 10 mg L^-1^ TZP stock, an 8:1 ratio of piperacillin: tazobactam (10 mg L^-1^ and 1.25 mg L^-1^, respectively) was used.

### p12 prevalence BLAST analysis

The plasmid p12 (11,744 bp, circular) was annotated with Bakta v1.9.4 (database v5.1), and replicon type and predicted mobility of the query were determined using MOB-suite^55^. Coding sequences and the replication origin (rep_origin), origin of transfer (oriT), and RNAI noncoding RNA were parsed from the resulting GenBank flat file using Biopython v1.87. The plasmid sequence was queried against the PLSDB plasmid database (v2024_05_31_v2; 72,556 records) using BLASTn (BLAST+ v2.17.0; blastn task, E-value ≤1×10^-10^, no percent-identity filter). Query coverage per subject (qcovs) and per-HSP alignment coordinates were retained for downstream analysis. Hits with qcovs ≥55 % were retained as informative matches, a threshold selected from a discontinuity in the coverage distribution across all 500 hits. Downstream data processing was performed in Python v3.13 using pandas v2.3.1, NumPy v2.3.2, and Matplotlib v3.10.5. The circular plasmid map was generated using Matplotlib’s polar plotting functions. The tool PlasmidFinder 2.0^56^ was used to determine the plasmid type, position, and replicon type of the other clinical plasmids used in this study.

### Plasmid transformation

KEIO wild-type and mutant strains (Δr*ecA*, Δ*recB*, Δ*recC*, and Δ*recD*) lacking the kanamycin cassette were transformed with plasmids of different sizes, including small plasmids in this study and corresponding larger plasmids generated previously by Heidarian *et al.,* 2026^27^. Competent cells of each strain were prepared by growing them overnight in 1 mL Difco^TM^ Luria-Bertani (LB) Lennox broth, and later subculturing in 50 mL LB, supplemented with 0.2% glucose, until OD_600_ = 0.3 - 0.4. Immediately, the cells were pelleted for 7 minutes at 4500 rpm at 4 °C, which was followed by three washes with ice-cold sterile dH_2_O. Plasmid purification was performed using the E.Z.N.A^®^ Plasmid DNA Mini Kit I (Omega Bio-Tek Inc.), according to the manufacturer’s instructions, and plasmid quality was assessed using the Nanodrop 1000 (Thermo Scientific). 50 μL of competent cells were mixed with 3 μL of pure plasmid and electroporated using the following settings: 2.5 kV, 200 Ω, 25 μF. The cells were immediately supplemented with LB + 0.2% glucose and left to recover at 37 °C. Transformants were plated on MHA + Ampicillin (AMP) in parallel with pure competent cells as a negative control. Colonies were grown overnight in the presence of AMP and saved at -80 °C in 10% dimethylsulfoxide (DMSO). Presence of the plasmid was confirmed with whole plasmid sequencing (Plasmidsaurus or Eurofins Genomics).

### PAP tests and HR determination

E-test data were first obtained to determine the average MIC of the population. Population analysis profile tests were performed using the following antibiotics: TZP, streptomycin (STR), and spectinomycin (SCM). Strains of interest were first isolated on MHA plates, and three independent overnight cultures were prepared by inoculating one colony in 1 mL MHB. The cultures were then diluted 1:1000 in PBS solution, and 1 μL of this dilution was used to inoculate 1 mL of fresh MHB. The grown bacterial cultures were diluted (10^-1^ to 10^-8^) in PBS buffer in a 96-well microtiter plate, as well as (1:20 to 1:2000) in 1.5 mL Eppendorf tubes, and plated in increasing antibiotic concentrations ranging from sub-MIC to multiple times above MIC. 5 μL drops of dilutions ranging from 10^-4^ to 10^-8^ were plated in technical triplicates on MHA plates containing the low concentrations of the antibiotic. For the higher antibiotic concentrations, 100 μL of dilutions 1:20 to 1:2000 were evenly spread with glass beads to avoid inoculum effects. Total viable counts were calculated by plating 100 μL of diluted (10^-6^ - 10^-7^) overnight cultures on MHA plates without antibiotics. Plates containing spots were incubated at 30 °C to ensure countable colonies after overnight incubation, and all other plates were incubated normally. Colonies were counted the next day, and plates were incubated again to allow for any slow-growing mutants to appear. The frequency of bacteria growing at the different antibiotic concentrations was calculated based on the total viable counts. Heteroresistance was noted when a bacterial subpopulation could grow at antibiotic concentrations ≥8-fold higher than the highest concentration the main susceptible population could withstand (HADNAG: Highest Amount of Drug Not Affecting Growth)^57^, at a frequency ≥10^-7^.

### Isolation of resistant mutants

For isolation of resistant mutants, the same protocol as above was followed, with the exception that bacteria were plated only on 2x, 4x, 8x, and 16x the HADNAG to select for resistance. The experiment was carried out in biological quadruplicates, on two independent days (n = 8). At least three individual colonies (or the maximum number of colonies on the plate), from each replicate and antibiotic concentration were re-streaked on MHA + TZP plates and grown overnight in MHB with the same TZP concentration, reaching a sample size ranging from n = 3 to n = 40. From the overnight growth, samples were saved at -20 °C for subsequent tests, including WGS, ddPCR, and E-tests. At the same time, each mutant was saved in 10% DMSO in an in-house strain collection at -80 °C.

### MIC determination

Previously saved samples of the resistant mutants were used for MIC analysis. Before freezing, the overnight growths were diluted 1:20 in PBS to reach 0.5 McFarland units, which corresponds to ±1.5×10^8^ CFU mL^-1^. The samples were thawed on ice, and each cell suspension was spread on MHA plates with a sterile cotton swab before placing an E-test strip (Liofilchem) in the middle and incubating the plates at 37 °C for ±20 hours as per the EUCAST guidelines.

### PCN determination by ddPCR

Plasmid copy number was determined using digital droplet PCR with primers targeting both the plasmid and the chromosome. For the chromosomal primers, either the lpp or rplT gene was used as a target, whereas for the plasmids, primers were designed to target regions outside the resistance cassettes. All oligonucleotide sequences are listed in Table S5. DNA was prepared by heat lysis of the cells. Briefly, pellets from overnight growths were resuspended in a 1:1 mixture of Fast Lysis Buffer (mericon DNA Bacteria kit, QIAGEN) and Nuclease-Free Water (Sigma-Aldrich). Samples were boiled for 10 minutes at 98 °C for cell lysis, followed by incubation at 12 °C to ensure DNA renaturation. Cell debris was removed by centrifugation, leaving pure DNA in the supernatant. The protocol used was based on a previously published study^58,59^, with a slight modification in the amount of restriction enzyme used, 0.5 μL instead of 1 μL (New England Biolabs). Droplet generation and reading were performed using the Bio-Rad QX200 system, PCR amplification using a C1000 Touch^TM^ Thermal Cycler (Bio-Rad), and data were analyzed using QuantaSoft Analysis Pro (v.1.0.569).

### Whole-genome sequencing through long- and short- reads, and sequence analysis

Some representative resistant mutants with high PCN from each parental strain (n = 19) were chosen for WGS. DNA preparation for both short- and long- read sequencing involved DNA extraction from pelleted cells using the MasterPure complete DNA&RNA purification kit (Epicentre), following the manufacturer’s instructions. DNA concentration was assessed using the Qubit 2.0 fluorometer and the dsBR Qubit assay kit (Invitrogen). Short-read sequencing using the DNBseq platform was performed by BGI (Warsaw, Poland) with paired-end libraries of ≤800 bp, and sequence analyses were done using the CLC Genomics Workbench (Qiagen). The sequences of the mutants were mapped against their corresponding parental, which was also sequenced and mapped against itself. We used the software’s own tools (Basic variant detection, InDel detection, Structural variant detection) to identify mutations, and then manually inspected both the chromosome and the plasmid to confirm their presence as well as identify IS element insertions, duplications, or other recombinations. A few representative revertant strains were also sequenced and analyzed in the same way.

Nanopore long-read sequencing was performed to assess the formation of plasmid multimers. Libraries were prepared according to the rapid barcoding kit 24 V14 protocol using the manufacturer’s conditions and sequenced using R10.4.1 flow cells on the MinION Mk1D instrument. For Nanopore, raw sequence reads were obtained in-house with base calling performed using MinKNOW v6.10.1 (Retrieved from ONT; https://github.com/nanoporetech/minknow_api) and Dorado v.2.0.1 (Retrieved from ONT; https://github.com/nanoporetech/dorado). The quality of reads was analyzed by NanoStat (v1.5.0)^60^.

For the analysis, a combined reference was constructed for each sample consisting of the host chromosome and the target plasmid. To enable continuous alignment of reads spanning tandem plasmid junctions, the plasmid sequence was duplicated head-to-tail within the reference. When additional plasmids were present, these were included as single-copy decoy contigs to minimize cross-plasmid mapping. Oxford Nanopore reads were aligned using minimap2 (v2.30; map-ont, --secondary=no)^61^, and alignments were sorted and indexed with SAMtools v1.22.1^62^. Primary and supplementary alignments were processed with Pysam (v0.22.1). Alignment coordinates were converted to read space, accounting for hard clipping and strand orientation, and overlapping intervals belonging to the same reference sequence were merged to avoid double-counting. For each read, two metrics were calculated: purity, defined as the proportion of the read aligned to a given reference sequence, and copy number, estimated as the cumulative aligned length on the target plasmid divided by the plasmid monomer length. Reads were classified as plasmid-derived when plasmid purity was ≥0.85 and further categorized as multimers (copy number ≥1.5) or monomers/fragments. Reads with chromosome or alternative plasmid purity ≥0.85 were assigned accordingly. Reads simultaneously exhibiting plasmid and chromosome (or alternative plasmid) purity ≥0.20 were classified as putative junction reads, whereas all remaining reads were designated unclassified or artifacts. The analysis pipeline was implemented in Python 3.12 using the standard library and Pysam.

Classification thresholds were evaluated by varying the purity cutoff between 0.50 and 1.00 and by re-examining a random subset of unclassified reads using sensitive remapping to distinguish genuine chromosome-plasmid junctions from chimeric or low-quality sequences. Because multimer detection requires reads to span at least 1.5 plasmid monomer lengths, the proportion of reads exceeding this minimum detectable length was calculated for each sample to account for differences in read-length distributions when comparing multimer frequencies.

To independently validate multimer assignments, a stratified subset of reads representing multimer, monomer, and negative-control classes was analyzed with cONcat^63^, an alignment-free method for reconstructing concatenated structures from Nanopore reads. Concordance between both approaches was assessed by comparing plasmid copy-number estimates and tandem-repeat structures. Figures were generated using Matplotlib^64^.

### Stability of HR phenotype

To determine how stable the HR phenotype was in mutants from both parentals, we started two individual bacterial cultures for each resistant mutant in 1.5 mL of MHB without antibiotics. The bacteria were passaged daily (1.5 μL in 1.5 mL fresh MHB) for 40 generations, assuming 10 generations of growth per day. After four days, the cultures were plated on MHA to isolate single clones. Three clones per plate per replicate (except for the mutants carrying the *recD* mutation, in which case 18 clones were isolated in total, from four biological replicates, to determine the plasmid loss rate caused by unsuccessful segregation of p12 concatemers) were isolated and grown overnight in plain MHB. An aliquot of each was stored at -80 °C in 10% DMSO, and more samples were saved at -20 °C for further tests, including WGS, E-tests, fitness analysis, and ddPCR.

### Growth rate determination

Growth rates of the parental strains, resistant mutants, and revertants were determined using the Bioscreen C apparatus (Oy Growth Curves Ab, Ltd). Six biological replicates were tested on separate days by inoculating thawed bacteria, straight from the -80 °C frozen stock, into fresh MHB media at a 1:1000 dilution. Three technical replicates (300 μL) were added to each well, and the plate was incubated for 24 hours at 37 °C inside the Bioscreen apparatus. Absorbance at 600 nm was measured every 4 minutes with shaking before each measurement. Analysis of the growth rates was performed using the BAT 2.1 software (Retrieved from https://thulin.shinyapps.io/bat2/), using the values obtained during exponential growth. An MG1655 strain was used as an internal control in every run, and all mutant growth rates were normalized to their respective parental strain, for which the growth rate was set to 1.

To assess the effect of carrying p12 on growth, we examined the growth rates of KEIO mutants Δ*recA*, Δ*recB*, Δ*recC*, Δ*recD*, and wild-type with and without p12 as described before, with the following exception. All strains were first streaked on MHA media, and overnight cultures were prepared by inoculating one colony in 1 mL of MHB. The cultures were then diluted 1:1000 in fresh media and inoculated in technical replicates in the Bioscreen plate. MG1655 was used again as an internal control and as the reference strain.

### Fitness change per kilobase of additional DNA

Since each resistant mutant contained different total DNA content, we wanted to determine the cost or benefit per kilobase of extra genetic material. To calculate this, we divided the fitness cost/advantage obtained by the BAT 2.1 software by the total amount of additional DNA in kb. This number represented the fitness change per kb. A positive value indicated a positive effect on growth rate by the acquisition of one kilobase of extra DNA, whereas a negative value represented growth impairment.

### RNA extraction

Samples were collected for RNA extraction as follows. Overnight cultures were prepared by inoculating a single colony in 1 mL MHB without any antibiotics (except in the case of TZP-resistant mutants, for which 1/2x MIC of TZP was added to ensure maintenance of the high copy number). Cultures were diluted 1:200 and grown in fresh media until mid-exponential phase. 4 mL of each culture was snap frozen in liquid nitrogen after the addition of 1 mL stop solution (95% ethanol, 5% phenol). The samples could be stored at -80 °C until needed or used immediately for RNA extraction.

RNA was extracted using a hot phenol-lysozyme protocol. Samples were thawed on ice and centrifuged at maximum speed for 10 minutes at 4 °C. The supernatant was discarded, and the pellets were resuspended in 600 μL of TE buffer (pH 8.0) supplemented with 0.5 mg mL^-1^ lysozyme. Immediately, 60 μL of 10% (w/v) SDS was added, and the samples were mixed by inversion and incubated in a 64 °C water bath for 1-2 minutes. 1:10 volume of 3 M NaOAc (pH 5.2) and 750 μL of acid phenol were added. Following vortexing, samples were incubated for 6 minutes at 64 °C and then on ice for 5 minutes. Samples were centrifuged at 13,000 rpm for 10 min at RT, and the aqueous phase was transferred to a new tube. One volume of chloroform was added, after which the samples were mixed and centrifuged under the same conditions. The aqueous phase was again transferred to a new tube, and RNA was precipitated by addition of 1.5 mL of an ice-cold 30:1 mixture of ethanol and 3 M NaOAc followed by incubation at -80 °C for 30 minutes. Samples were then centrifuged at 13,000 rpm for 30 minutes at 4 °C, and the pellets were rinsed with ice-cold 80% ethanol. Residual ethanol was removed by briefly centrifuging the samples, carefully aspirating any remaining liquid, and repeating the wash twice. The pellets were dried at RT for a couple of minutes, and RNA was resuspended in 40 μL sterile dH_2_O by shaking at 900 rpm and 65 °C for 3 minutes. RNA concentration and quality were assessed by using the Nanodrop 1000 (Thermo Scientific) and gel electrophoresis on a 1% agarose gel, respectively.

To remove genomic DNA contamination, the RNA samples were DNase-treated using the TURBO DNA-free^TM^ kit (Invitrogen). Approximately 35 μg of RNA, 1 μL of TURBO DNase enzyme, 1x of DNase buffer (10x), and water were mixed and incubated for 20 minutes at 37 °C, to activate the enzyme. The RNAs were purified and reprecipitated using a phenol-chloroform-isoamyl alcohol (PCI) based protocol. For this, an equal volume of PCI was added to each sample, which was then centrifuged to separate the aqueous phase containing the RNA from the oil phase. Precipitation continued as mentioned previously, and the quality of the RNA was assessed again.

### Northern blot

RNA samples were loaded on a 6% polyacrylamide gel, containing 8 M urea, 1x TBE, APS, and TEMED. 10 μg of RNA were mixed with 2x Gel Loading Buffer II (GLII) to a final volume of 20 μL. Samples were denatured for 4 minutes at 95 °C and cooled down on ice for another 5 minutes. A radioactive pUC19 MSP1 marker was used as a ladder. Gel electrophoresis was conducted for approximately 1.5 hours at 20 W, ∼ 420 V. A wet transfer of the gel to a nitrocellulose membrane (Hybond-XL, GE Healthcare) was performed for 2 hours at 6 W, 340 mA at 4 °C. Following UV cross-linking at 1200 mJ/cm², the membranes were incubated in Church buffer (0.25 M sodium phosphate buffer (pH 7.2), 1 mM EDTA, and 0.07% SDS) for 45 min at 42 °C to block nonspecific binding sites, and subsequently, [γ-^32^P] ATP 5′-end-labeled DNA probes targeting RNAI, RNAII and 5S rRNA as a loading control (listed in Table S5) were denatured and added one by one to initiate hybridization, which was carried out overnight at 42 °C. After hybridization, the membrane was washed twice with 2x SSC/0.1% SDS buffer, dried, sealed in plastic wrap, and exposed to a phosphorscreen for 2-3 days. Radioactive signals were visualized using a Typhoon FLA 7000 phosphorimager (GE Healthcare). The membrane was then stripped by placing it in a boiling 0.1% SSC/0.1% SDS buffer and subsequently pre-hybridized for the next probe.

### Multimer separation by gel electrophoresis

Plasmid DNA was isolated using the NucleoBondTM Xtra Midi kit (MACHEREY-NAGEL) according to the manufacturer’s instructions, and DNA concentration was measured with the Qubit 2.0. p12 DNA was digested with HindIII, following standard protocols from the manufacturer (New England Biolabs). For each gel, the same amount of DNA was loaded into each well after adjusting the DNA concentration based on the Qubit measurements. To ensure proper separation, a 0.4% agarose gel in 1x TAE was prepared, and the gels were run at 3 V/cm for 16-18 hours. For multimer detection of high-molecular-weight fragments, the GeneRuler High Range DNA Ladder (ThermoFischer) was used, whereas for smaller fragments, the GeneRuler 1 kb DNA ladder (ThermoFischer) was used.

### Statistical analyses

Most statistical tests were performed in GraphPad Prism 11, and to choose the appropriate test for each dataset, the normality/lognormality of the data was first assessed by a Shapiro-Wilk test, the standard for small sample sizes. The linear regression models were performed in RStudio, and all tests from Nanopore long-read sequencing analysis were done with Python.

## FIGURES

**Supplementary Figure 1.**
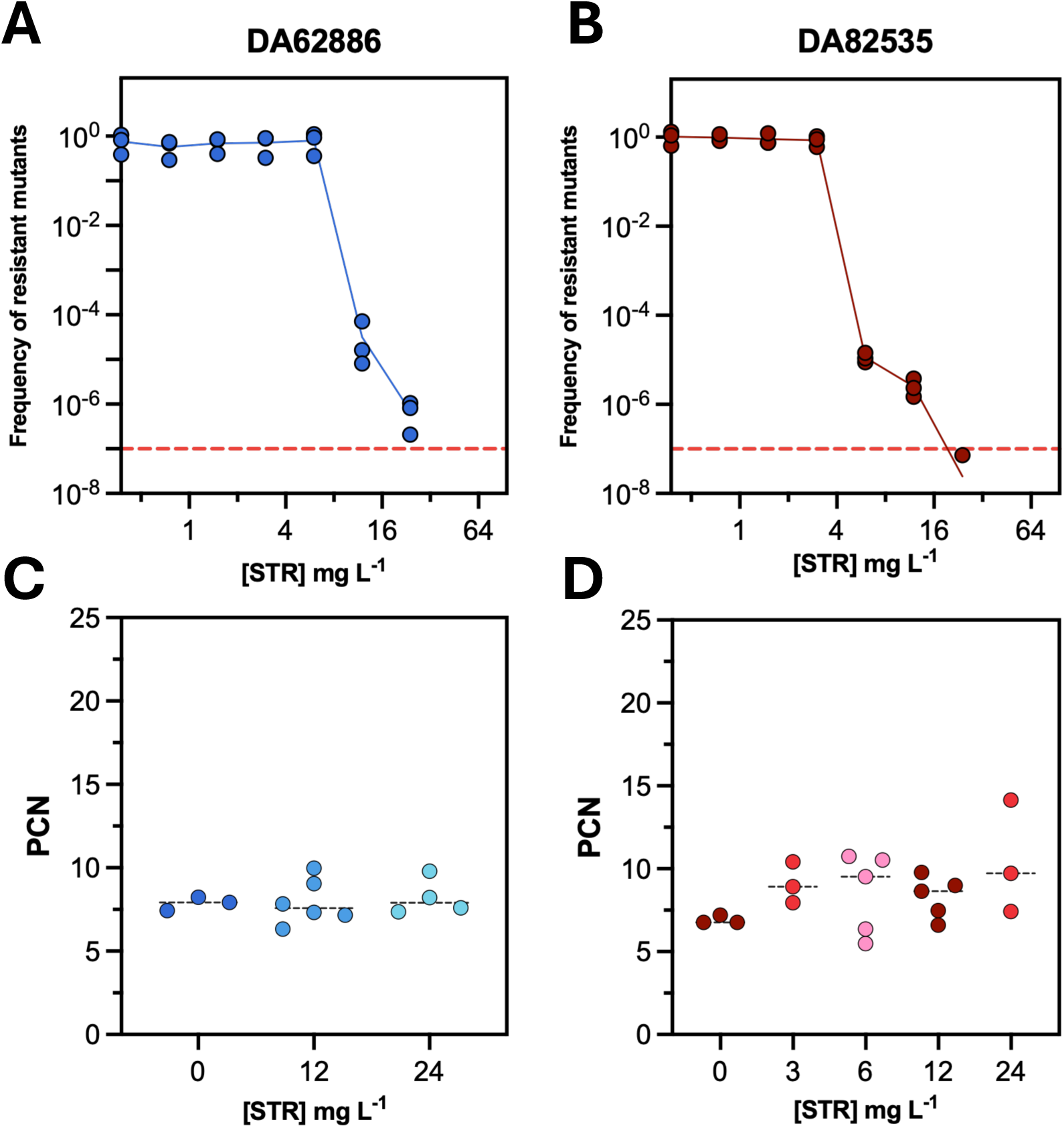
No HR was observed for STR in the p12-carrying strains. **(A-B).** PAP tests for STR of DA62886 and DA82535, respectively, carrying p12. Dashed red lines represent the frequency threshold (≥10^-7^) (n = 3). **(C-D).** PCN of isolated resistant mutants at (G) 2x, and 4x MIC of the main susceptible population for DA62886 and (H) 1x, 2x, 4x, and 8x MIC for DA82535. Data are presented as the mean of all replicates. Two-tailed Mann-Whitney tests were performed to compare all groups with the control (0 mg L^-1^ STR).

**Supplementary Figure 2.**
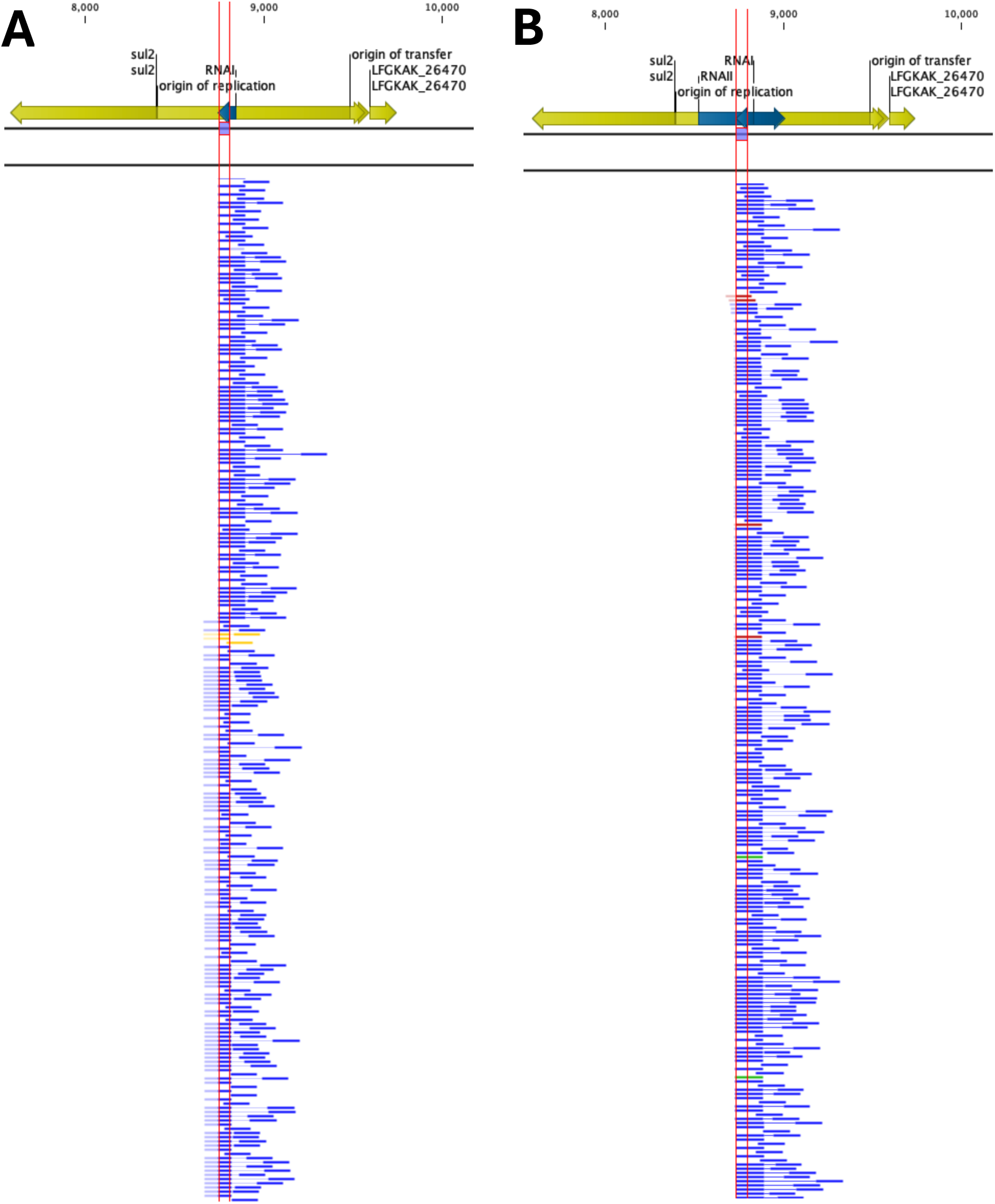
The 65-bp duplication located in the p12 RNAI/RNAII region can only be found in a subset of the population. Representative images of WGS analysis, for **(A)** a 65-duplication resistant mutant and **(B)** a corresponding revertant, revealing that the mutation was not present in all reads, and the proportion was particularly lower in the revertant, with barely any reads showing the mutation. More information can be found in Table S4 of the Supplementary Material.

**Supplementary Figure 3.**
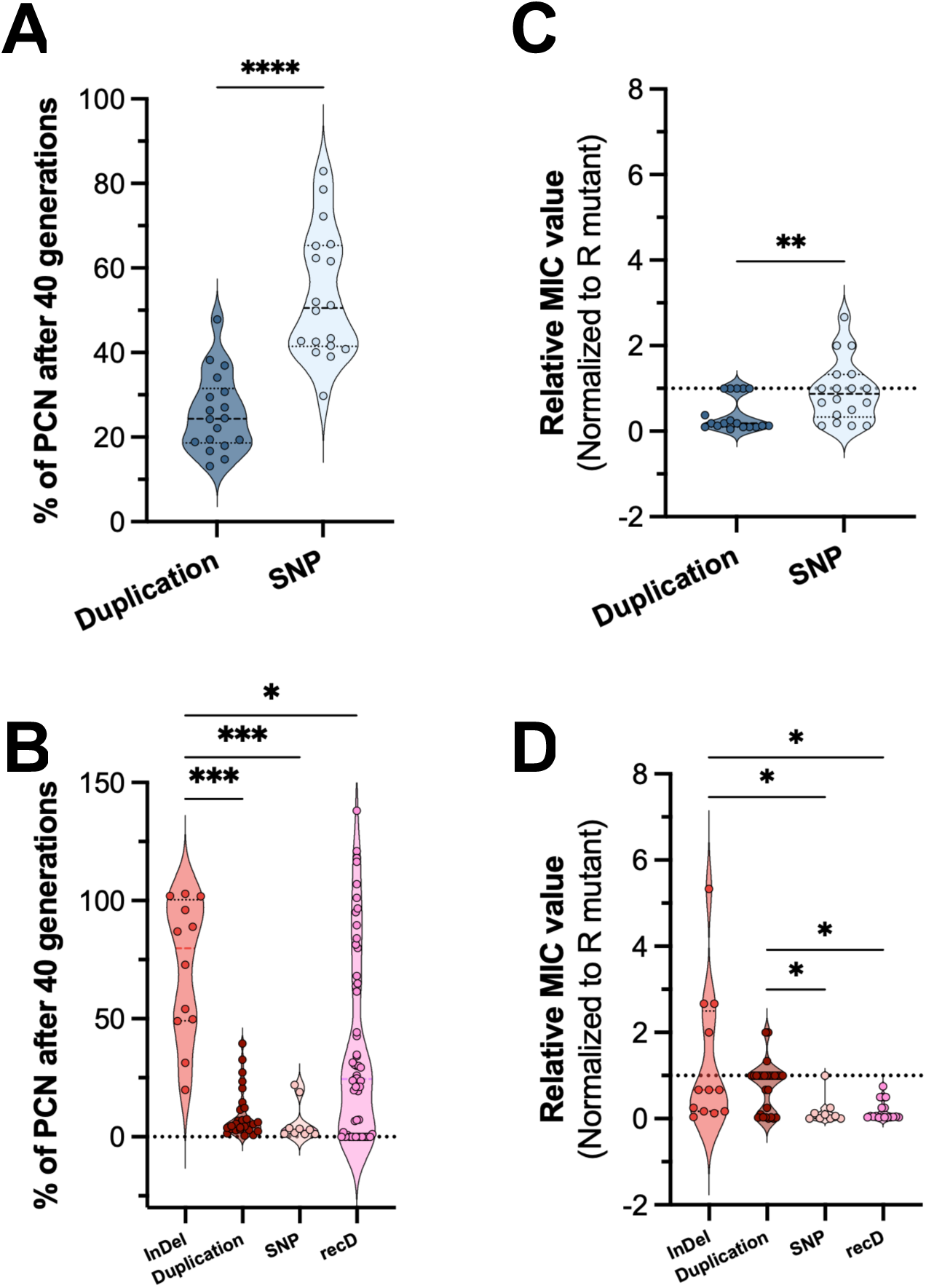
The rate of reversion of increased PCN depends on the mutations present. **(A-B).** Percentage of the initial high PCN remaining on the revertants after 40 generations of growth without selection, for DA62886 and DA82535 mutant backgrounds, respectively. Resistant mutants harboring the same mutation type were grouped, and individual colonies of each were analyzed. An unpaired two-tailed t-test was performed for DA62886 and Kruskal-Wallis with Dunn’s correction for DA82535 (based on the Shapiro-Wilk normality test). **(C-D).** Relative MIC_TZP_ values of the revertants, normalized to the initial MIC value of the resistant mutant. Two-tailed Mann-Whitney and Kruskal-Wallis with Dunn’s post-hoc correction were performed, respectively. *: p<0.05, **: p<0.01, ***: p<0.001, ****: p<0.0001

**Supplementary Figure 4.**
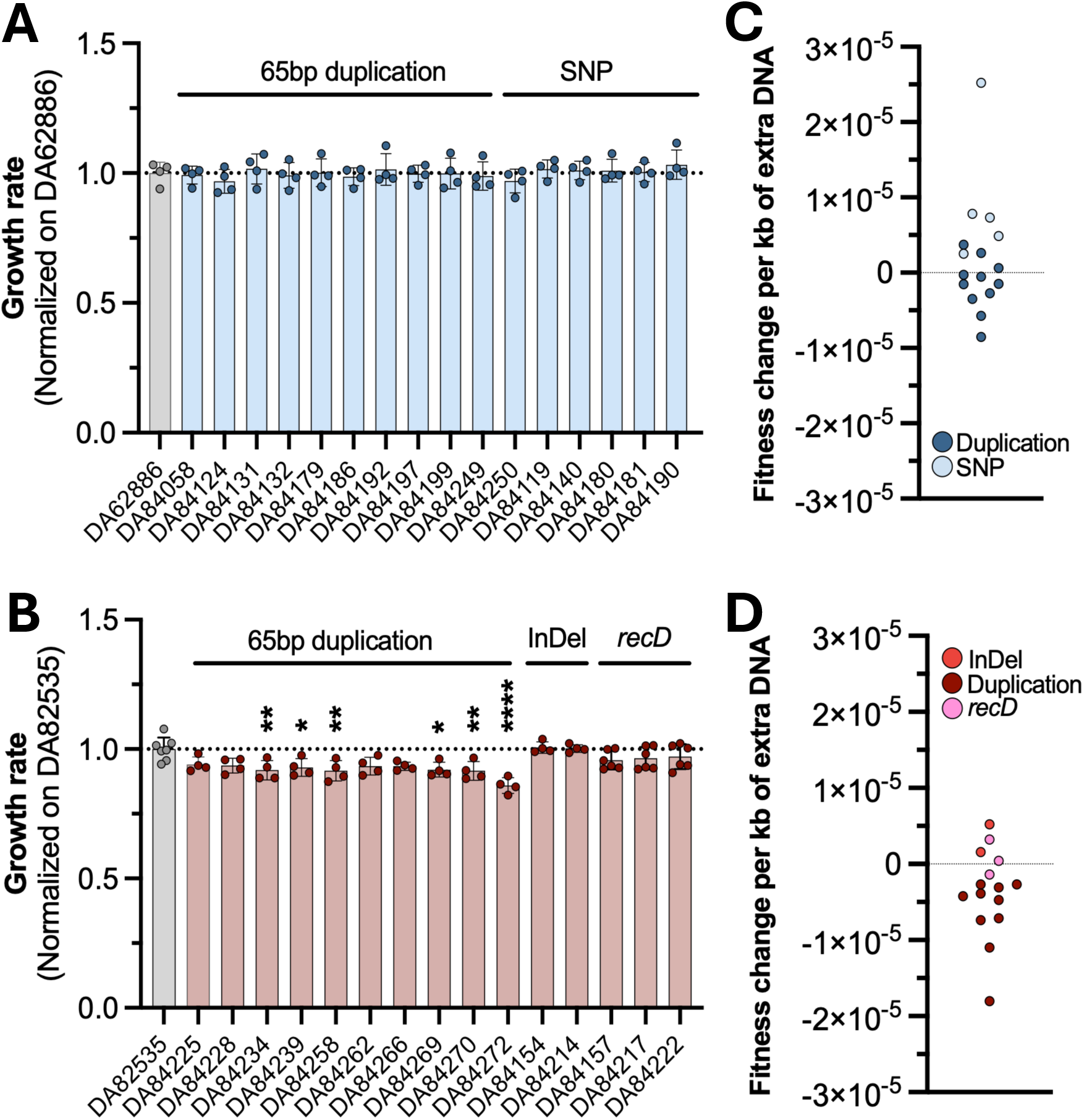
Fitness cost does not strongly correlate with the amount of extra DNA. **(A-B).** Growth rates of TZP-resistant mutants carrying only one mutation type. Data are shown as mean ± SD (n = 4 for all except n = 7 for DA82535), and each sample is normalized to the parental mean growth rate. The dotted line represents the reference growth rate of the parental, at 1. Normality was assessed using a Shapiro-Wilk test, and a parametric one-way ANOVA (with Dunnett’s correction) was performed to compare each mutant to the parental (*: p<0.05, **: p<0.01, ***: p<0.001, ****: p<0.0001). **(C-D).** Effect of carrying additional DNA on fitness. The amount of additional DNA was calculated for each mutant based on PCN values, and the fitness change of carrying one kb of extra DNA on bacterial fitness was calculated based on the relative growth rates obtained from (A-B).

**Supplementary Figure 5.**
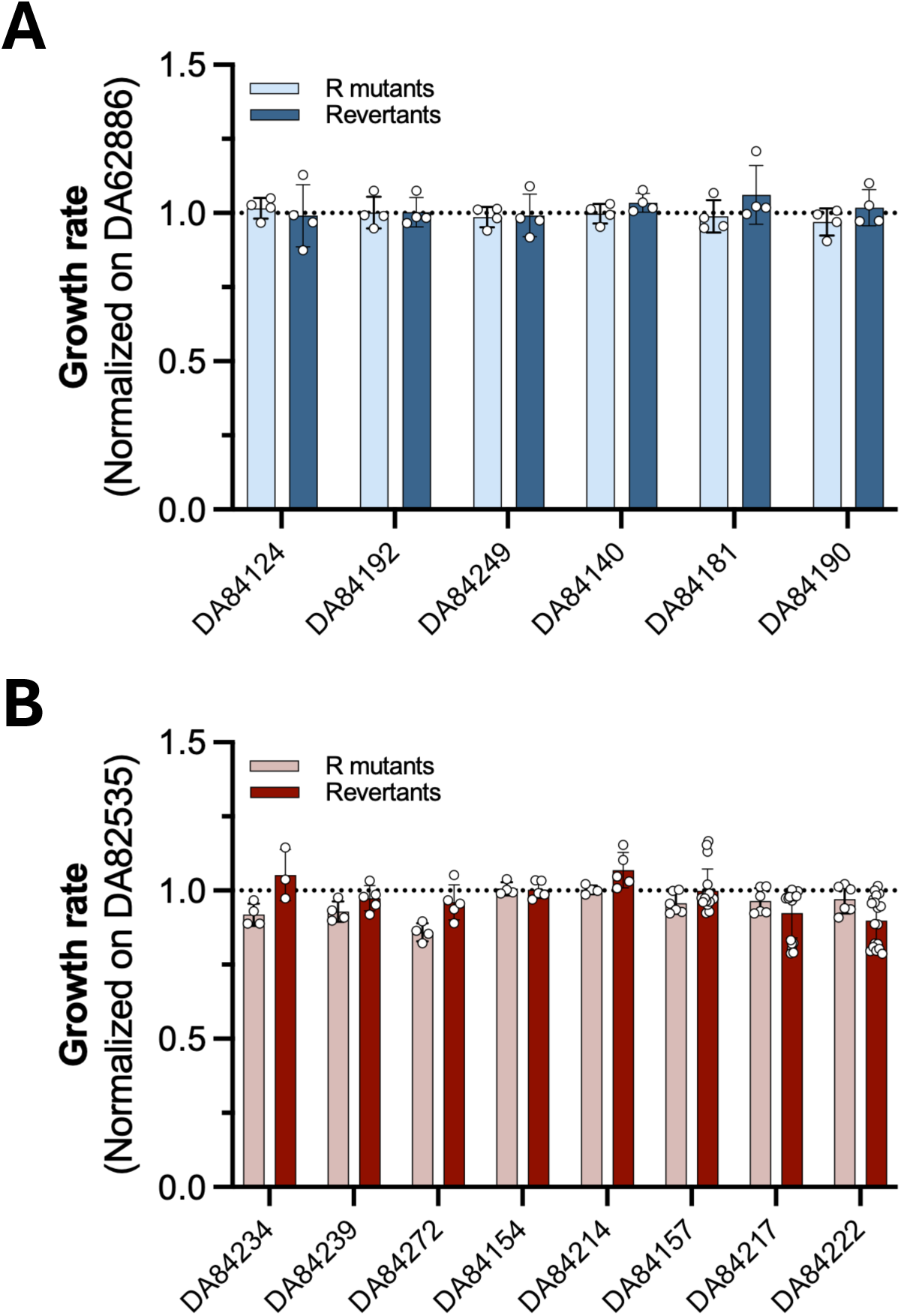
Growth rates of mutants determined after 40 generations of growth without selection. **(A-B).** Growth rates of TZP-resistant mutants and revertants after 40 generations of growth with no selection. Data are shown as mean ± SD of four independent measures of fitness for the resistant mutants and three to 18 measures of independent revertant strains. Normality was assessed using a Shapiro-Wilk test, and a parametric one-way ANOVA was performed to compare the revertants to the resistant mutant.

**Supplementary Figure 6.**
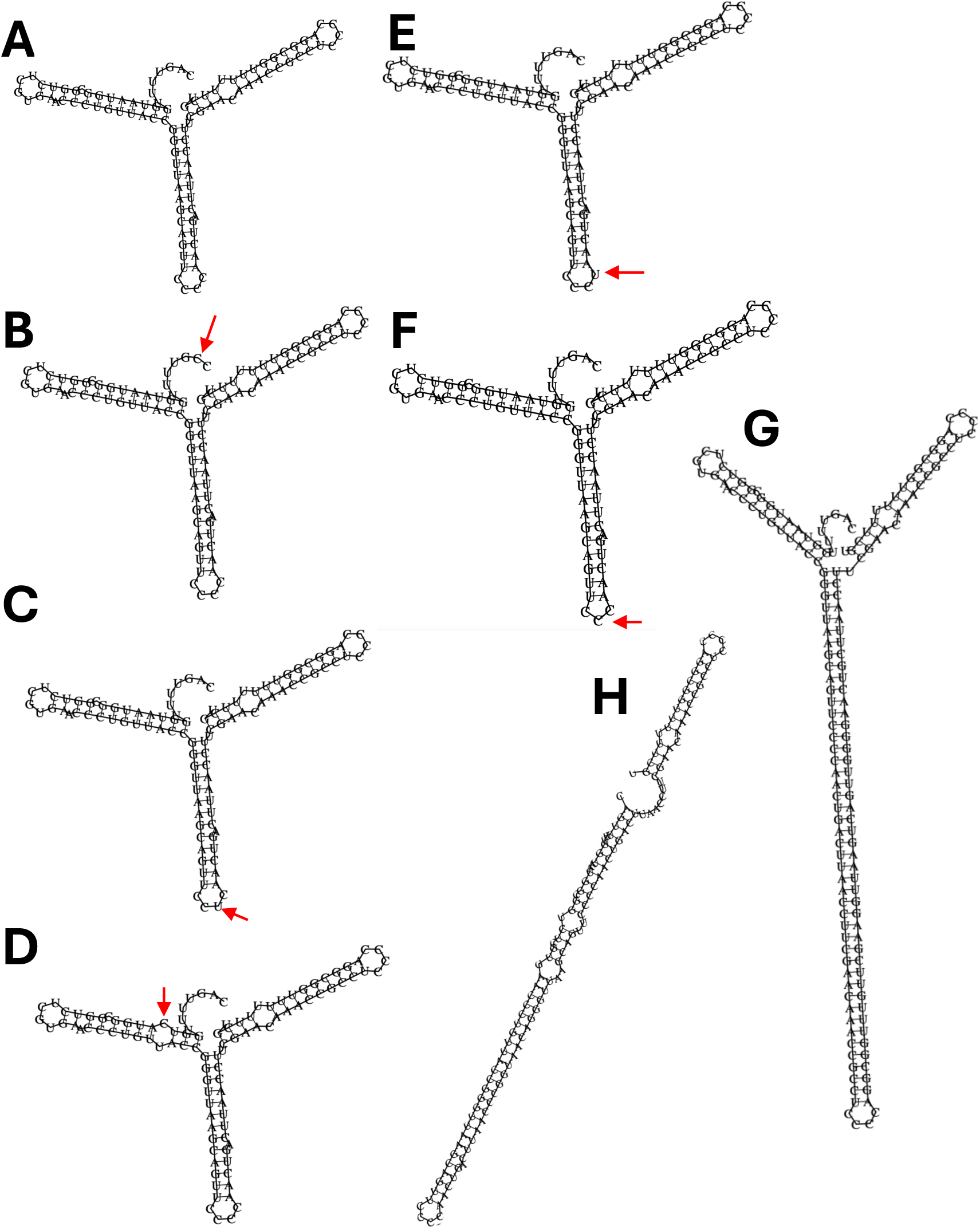
Predicted RNAI secondary structure. All predictions were made using the RNAfold server (Retrieved from http://rna.tbi.univie.ac.at/cgi-bin/RNAWebSuite/RNAfold.cgi). Red arrows represent the position of the mutation **(A)**. Parental p12 RNAI **(B-E).** p12 RNAI harboring a SNP at different locations **(F).** p12 RNAI harboring an InDel **(G).** p12 RNAI with a 65 bp duplication **(H).** p12 RNAI with a 38 bp duplication.

**Supplementary Figure 7.**
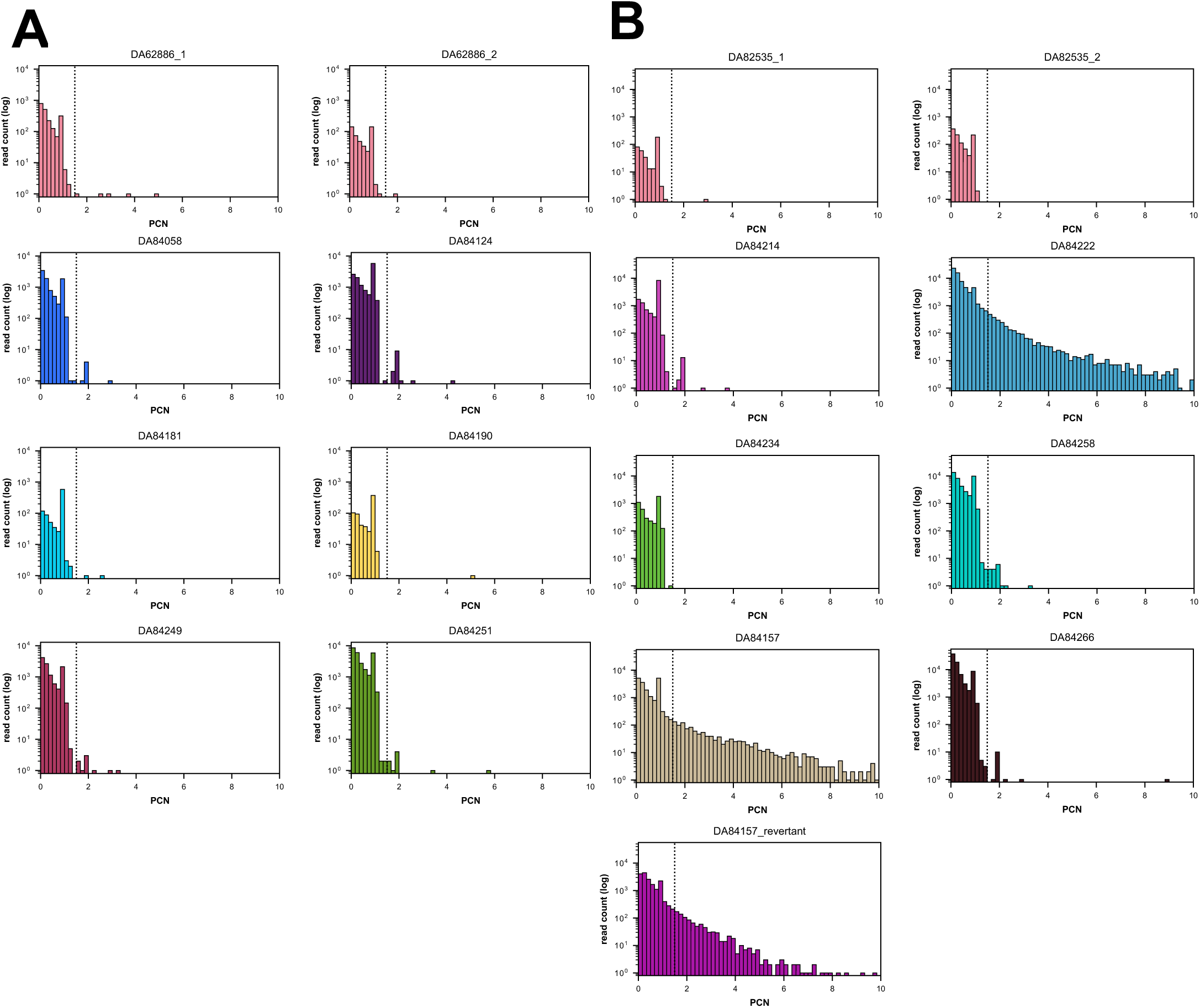
Copy-number distributions of plasmid-derived reads across high-PCN mutants. **(A).** DA62886 chromosome background; n = 8 samples: the parental strain sampled before and after selection (labelled “1” and ”2”, respectively) and six individually sequenced PCN mutant isolates. **(B)** DA82535 chromosome background; n = 9 samples: the parental strain before and after selection (“1”/”2”), six individual PCN mutant isolates, and one antibiotic-revertant isolate. For each sample, the histogram shows PCN, the estimated plasmid copy number spanned by a read, computed only among reads already passing the purity_plasmid ≥ 0.85 threshold, plotted on a log-scaled y-axis. The dotted vertical line marks the multimer threshold (PCN ≥ 1.5) used to call a read a plasmid multimer rather than a monomer or fragment.

**Supplementary Figure 8.**
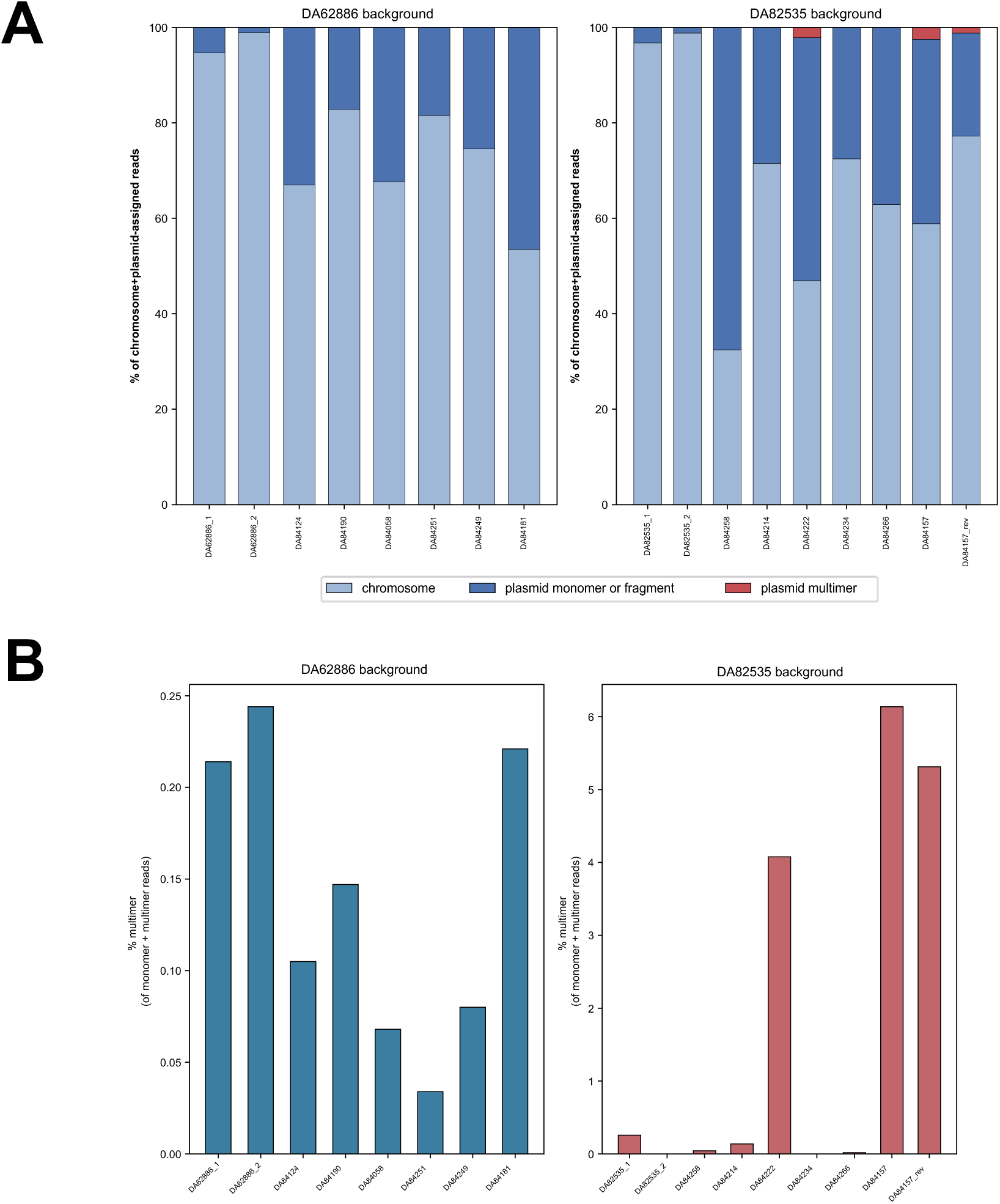
Read classification composition and multimer percentage across the high-PCN mutants, split by chromosome background. **(A).** Stacked bar chart of read-class composition (% of chromosome + plasmid-assigned reads) for each sample, split into i) DA62886 background and ii) DA82535 background; bars are colored by read class: chromosome, plasmid monomer/fragment, plasmid multimer, as shown in the legend. **(B)** Bar chart of % multimer of monomer + multimer reads for the same 17 samples, split the same way; each bar represents a single measurement (n = 1 per sample).

**Supplementary Figure 9.**
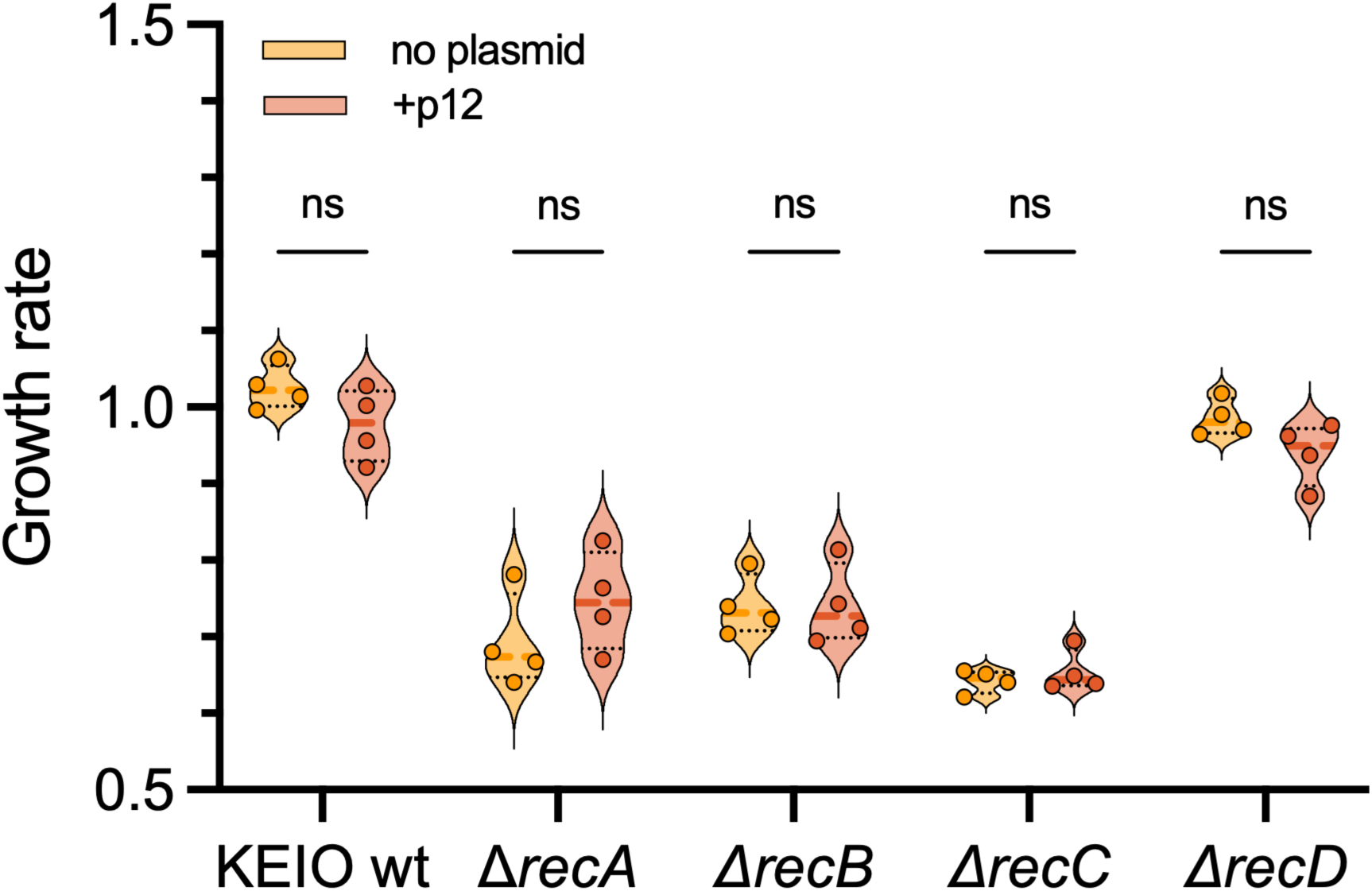
Carrying p12 does not confer a fitness cost for the host. Growth rates of KEIO wild-type and recombination mutants with and without p12. Data are shown as mean ± SD of 4 independent biological replicates. Normality was assessed using a Shapiro-Wilk test and a parametric one-way ANOVA was performed to compare all the groups (ns: non-significant).

**Supplementary Figure 10.**
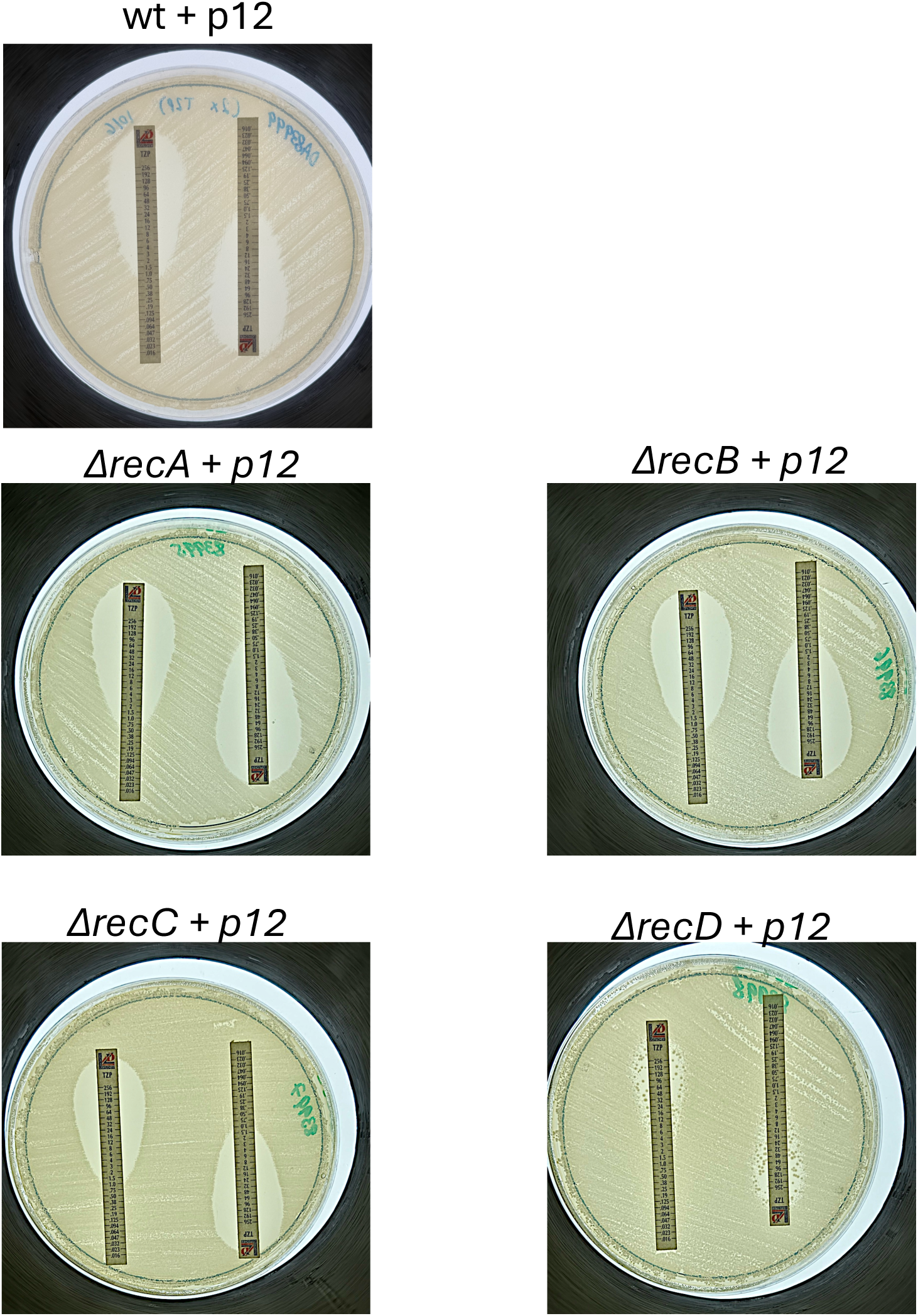
A Δ*recD +* p12 strain exhibits an HR-like phenotype on an E-test. E-tests for TZP were performed in technical duplicates for **(A).** KEIO wild-type (wt) + p12, **(B).** Δ*recA* + p12, **(C).** Δ*recB* + p12, **(D).** Δ*recC* + p12, and **(E).** Δ*recD* + p12

**Supplementary Figure 11.**
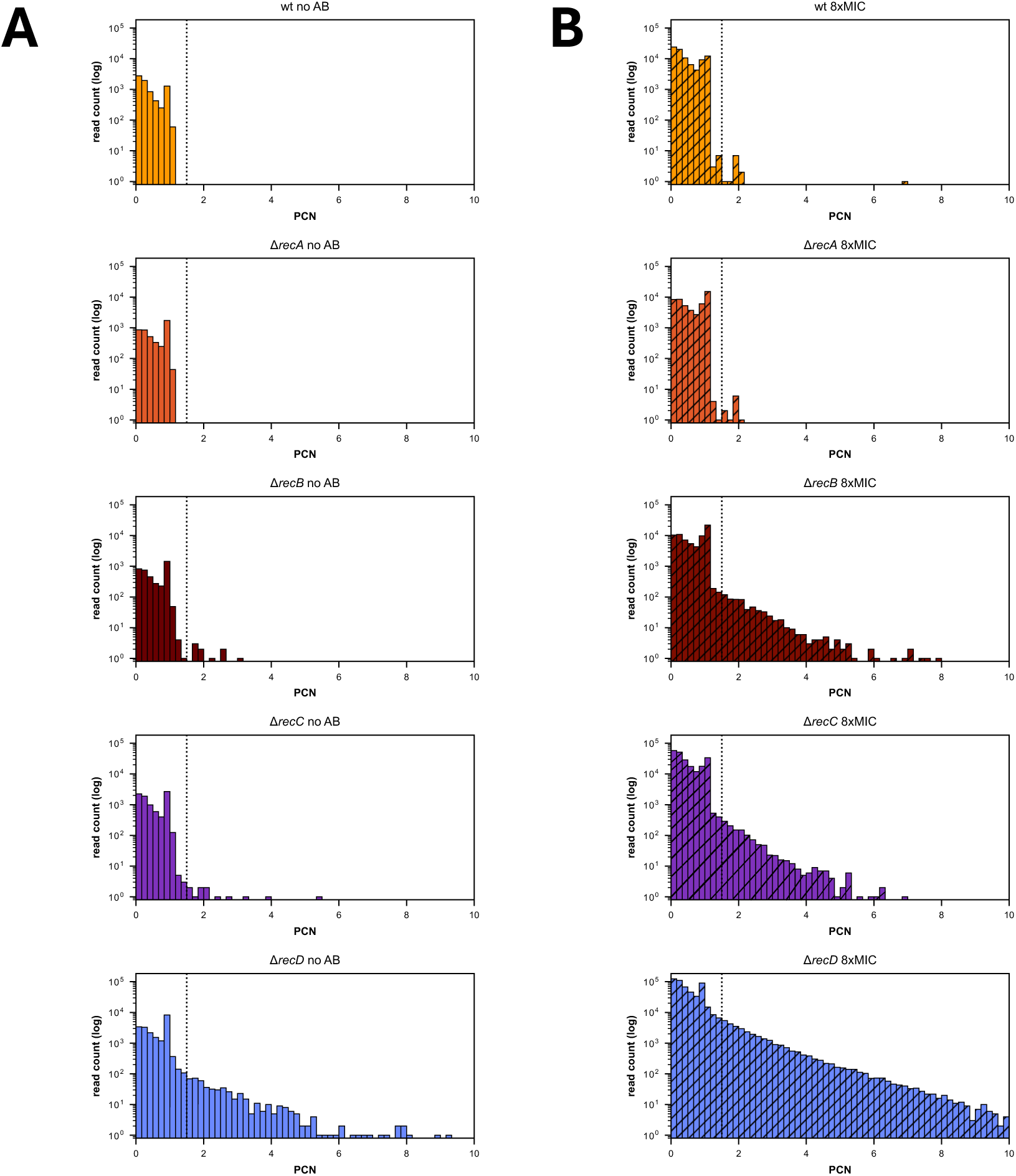
Distribution of plasmid copy number estimates across RecA-RecBCD mutants. Histograms show the distribution of plasmid copy number estimates (PCN) for individual plasmid-derived reads (plasmid purity ≥ 0.85) from wild-type (wt), Δ*recA*, Δ*recB*, Δ*recC*, and Δ*recD* strains grown **(A)** in the absence of antibiotic (no AB) or **(B)** following selection at 8x MIC. The vertical dashed line indicates the threshold (PCN = 1.5) used to distinguish monomeric from multimeric plasmid molecules, with reads to the right of the threshold classified as putative multimers. Histograms are displayed using a common x-axis scale (0–10) to facilitate comparison across genotypes and conditions.

**Supplementary Figure 12.**
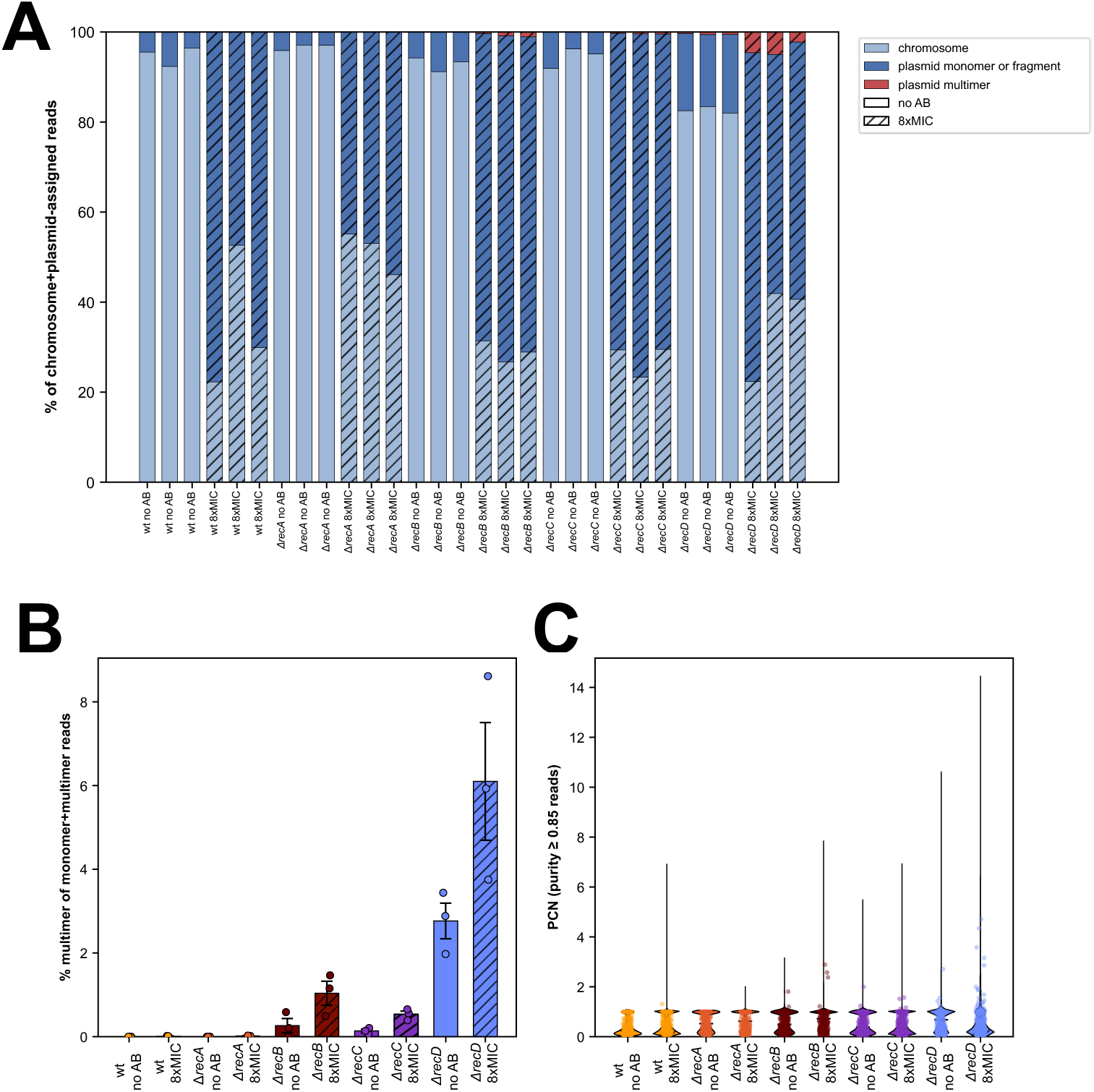
Effect of RecA-RecBCD pathway mutations on plasmid multimerization and plasmid copy number. **(A).** Distribution of read classification categories across all RecA-RecBCD mutants. Stacked bar plots show the percentage of sequencing reads assigned to chromosomal, plasmid monomer or fragment, plasmid multimer, junction candidate, or unclassified/artefactual categories for wild-type (wt), Δ*recA*, Δ*recB*, Δ*recC*, and Δ*recD* strains grown in the absence of antibiotic (no AB) or following selection at 8xMIC (represented by diagonal stripes). Percentages are calculated relative to the total number of reads per sample, with each bar representing one biological replicate. **(B).** Plasmid multimerization, expressed as the percentage of multimeric molecules among confidently classified plasmid molecules (monomer + multimer), for wild-type (wt), Δ*recA*, Δ*recB*, Δ*recC*, and Δ*recD* strains grown in the absence of antibiotic (no AB) or following selection at 8xMIC. Each point represents an independent biological replicate (n = 3). Boxes indicate the interquartile range (IQR), the center line denotes the median, and whiskers extend to 1.5 x IQR. Colors indicate genotype, and diagonal stripes represent antibiotic-selected samples. **(C)**. Distribution of plasmid copy number estimates (PCN) for individual plasmid-derived reads with plasmid purity ≥ 0.85. Violin plots show the distribution of all reads passing the purity threshold, with overlaid points representing individual reads.

**Supplementary Figure 13.**
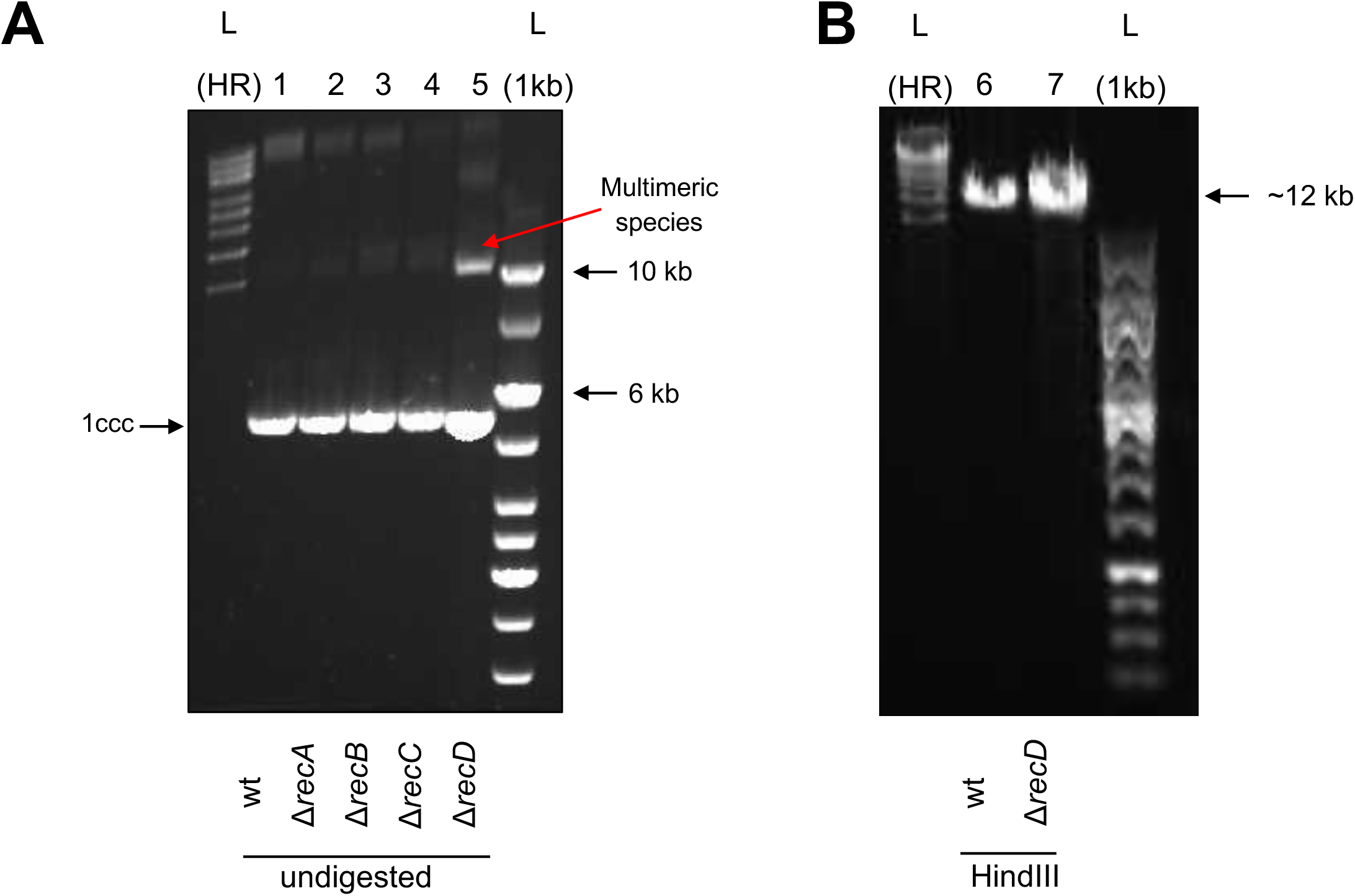
*recD* mutants exhibit increased p12 multimer formation. **(A).** Extracted undigested plasmid DNA (p12) from strains: KEIO wild-type, Δ*recA*, Δ*recB*, Δ*recC*, and Δ*recD* (lanes 1-5) and **(B)** digested p12 of KEIO wild-type and Δ*recD* with the 1-cutter restriction enzyme, HindIII (lanes 6 and 7). High-range DNA ladder (left), and a 1 kb DNA ladder (right). 1ccc: covalently closed circular monomer

**Supplementary Figure 14.**
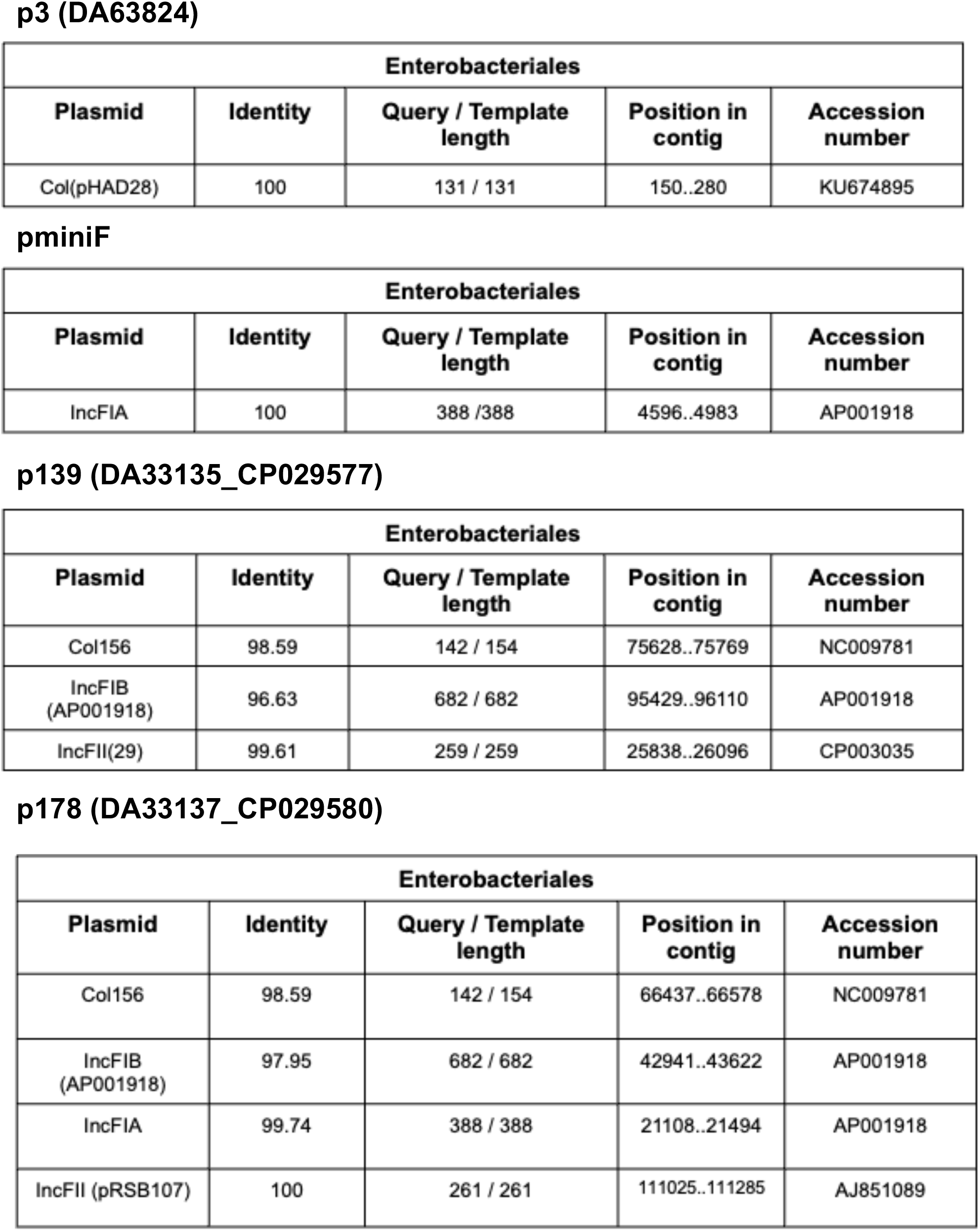
Bioinformatic analysis reveals the replicon type of the plasmids. Use of PlasmidFinder 2.0 to identify the replicons for different clinical plasmids used in this study.

**Supplementary Figure 15.**
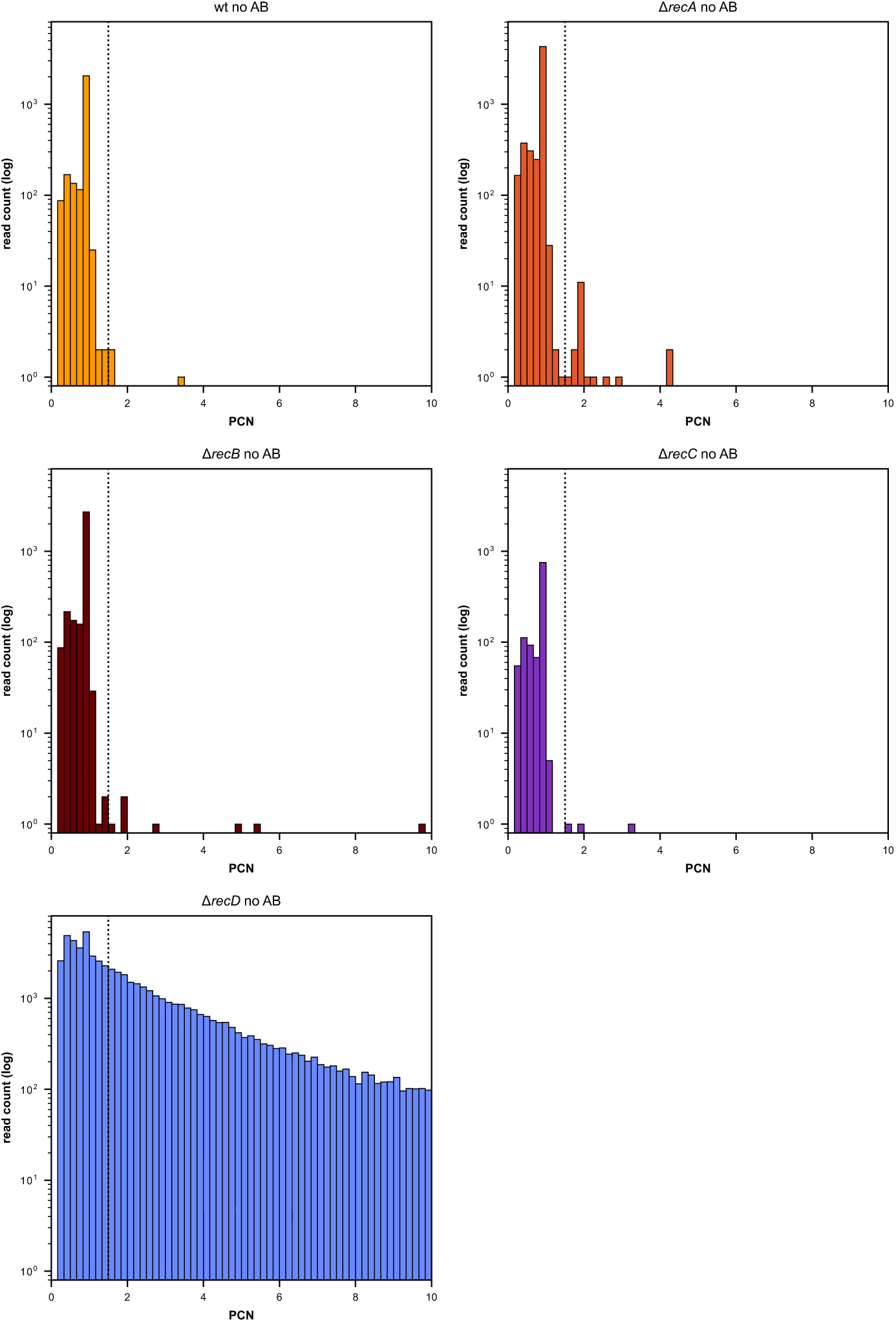
Distribution of plasmid copy number estimates across RecA-RecBCD mutants in p3-carrying strains. Histograms show the distribution of plasmid copy number estimates (PCN) of the p3 plasmid for individual plasmid-derived reads (plasmid purity ≥ 0.85) from wild-type (wt), Δ*recA*, Δ*recB*, Δ*recC*, and Δ*recD* strains. The vertical dashed line indicates the threshold (PCN= 1.5) used to distinguish monomeric from multimeric plasmid molecules, with reads to the right of the threshold classified as putative multimers. Histograms are displayed using a common x-axis scale (0–10) to facilitate comparison across genotypes.

**Supplementary Figure 16.**
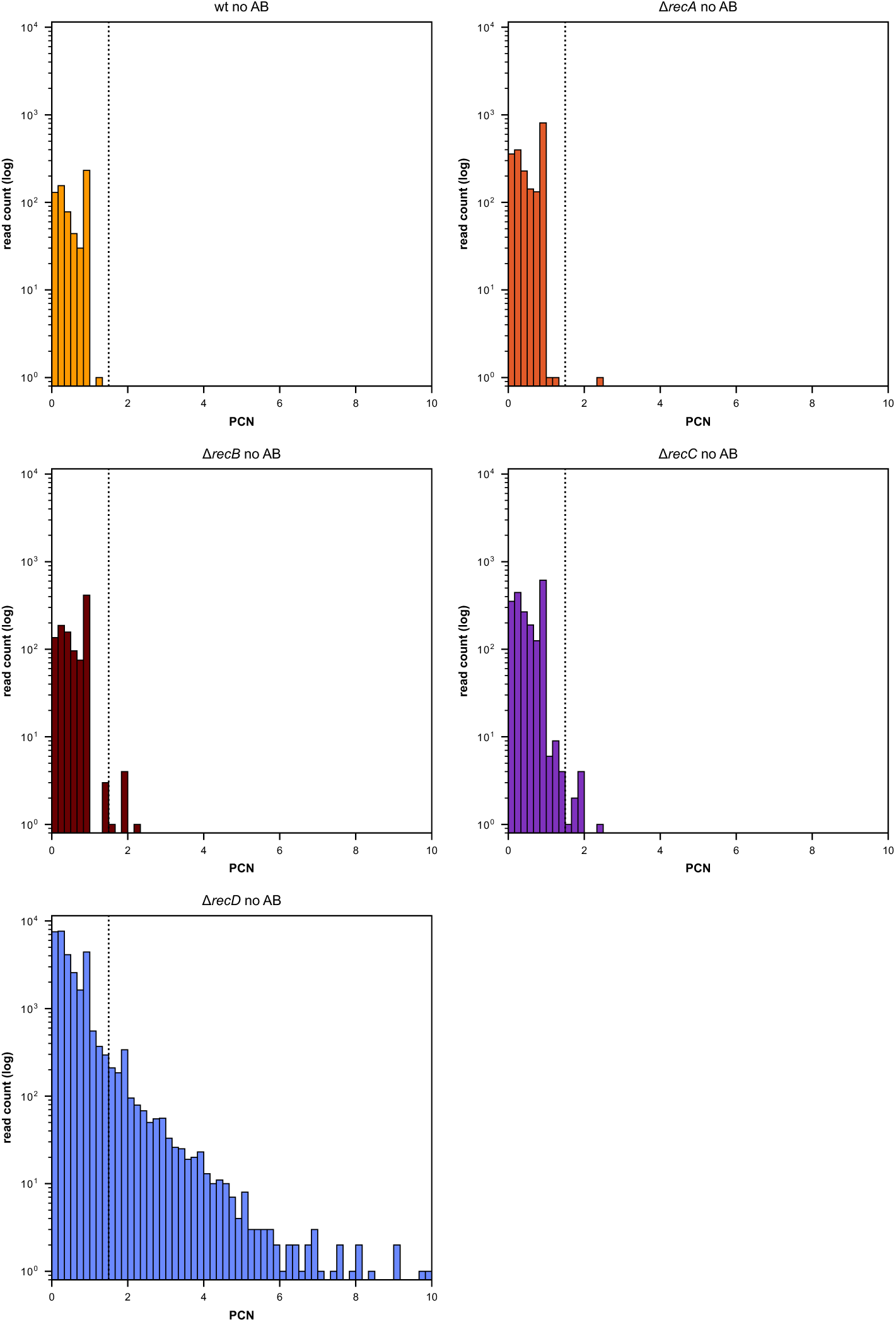
Distribution of plasmid copy number estimates across RecA-RecBCD mutants in miniF-carrying strains. Histograms show the distribution of plasmid copy number estimates (PCN) of the miniF **plasmid** for individual plasmid-derived reads (plasmid purity ≥ 0.85) from wild-type (wt), Δ*recA*, Δ*recB*, Δ*recC*, and Δ*recD* strains. The vertical dashed line indicates the threshold (PCN = 1.5) used to distinguish monomeric from multimeric plasmid molecules, with reads to the right of the threshold classified as putative multimers. Histograms are displayed using a common x-axis scale (0–10) to facilitate comparison across genotypes.

**Table S1.** List of bacterial strains used in this study.

| Strain ID | Description | Genotype | + Plasmid | Reference |
| --- | --- | --- | --- | --- |
| DA62886 | BSI clinical isolate |  | p12 (11,744 bp) | Heyman et al., 2025 |
| DA82535 | MG1655 lab strain |  | p12 (11,744 bp) | This study |
| DA83999 | KEIO wild-type | Eco wt ME9062 BW25113 *rrnB*<br>ΔlacZ4787 *hsdR*514 $\Delta$(araBAD)567<br>$\Delta$(rhaBAD)568 *rph*-1" | p12 (11,744 bp) | This study |
| DA83995 | *recA* deletion | F-, $\Delta$(araD-araB)567,<br>ΔlacZ4787(::rrnB-3), λ-, ΔrecA774,<br>*rph*-1, $\Delta$(rhaD-rhaB)568, *hsdR*514 | p12 (11,744 bp) | This study |
| DA83996 | *recB* deletion | F-, $\Delta$(araD-araB)567,<br>ΔlacZ4787(::rrnB-3), λ-, ΔrecB745,<br>*rph*-1, $\Delta$(rhaD-rhaB)568, *hsdR*514 | p12 (11,744 bp) | This study |
| DA83997 | *recC* deletion | F-, $\Delta$(araD-araB)567,<br>ΔlacZ4787(::rrnB-3), λ-, ΔrecC747,<br>*rph*-1, $\Delta$(rhaD-rhaB)568, *hsdR*514 | p12 (11,744 bp) | This study |
| DA83998 | *recD* deletion | F-, $\Delta$(araD-araB)567,<br>ΔlacZ4787(::rrnB-3), λ-, ΔrecD744,<br>*rph*-1, $\Delta$(rhaD-rhaB)568, *hsdR*514 | p12 (11,744 bp) | This study |
| DA63824 | BSI clinical isolate |  | p3 (2,699 bp) | Heyman et al., 2025 |
| DA84000 | KEIO wild-type | Eco wt ME9062 BW25113 *rrnB*<br>ΔlacZ4787 *hsdR*514 $\Delta$(araBAD)567<br>$\Delta$(rhaBAD)568 *rph*-1" | p3 (2,699 bp) | This study |
| DA84001 | *recA* deletion | F-, $\Delta$(araD-araB)567,<br>ΔlacZ4787(::rrnB-3), λ-, ΔrecA774,<br>*rph*-1, $\Delta$(rhaD-rhaB)568, *hsdR*514 | p3 (2,699 bp) | This study |
| DA84002 | *recB* deletion | F-, $\Delta$(araD-araB)567,<br>ΔlacZ4787(::rrnB-3), λ-, ΔrecB745,<br>*rph*-1, $\Delta$(rhaD-rhaB)568, *hsdR*514 | p3 (2,699 bp) | This study |
| DA84003 | *recC* deletion | F-, $\Delta$(araD-araB)567,<br>ΔlacZ4787(::rrnB-3), λ-, ΔrecC747,<br>*rph*-1, $\Delta$(rhaD-rhaB)568, *hsdR*514 | p3 (2,699 bp) | This study |
| DA84004 | *recD* deletion | F-, $\Delta$(araD-araB)567,<br>ΔlacZ4787(::rrnB-3), λ-, ΔrecD744,<br>*rph*-1, $\Delta$(rhaD-rhaB)568, *hsdR*514 | p3 (2,699 bp) | This study |
| DA81984 | KEIO wild-type | Eco wt ME9062 BW25113 *rrnB*<br>ΔlacZ4787 *hsdR*514 $\Delta$(araBAD)567<br>$\Delta$(rhaBAD)568 *rph*-1" | p139 (139,191 bp) | Heidarian et al., 2026 |
| DA81965 | *recA* deletion | F-, $\Delta$(araD-araB)567,<br>ΔlacZ4787(::rrnB-3), λ-, ΔrecA774,<br>*rph*-1, $\Delta$(rhaD-rhaB)568, *hsdR*514 | p139 (139,191 bp) | Heidarian et al., 2026 |
| DA81966 | *recB* deletion | F-, $\Delta(\text{araD-araB})567$,<br>$\Delta\text{lacZ4787}(\text{::rrnB-3})$, $\lambda^-$, $\Delta\text{recB745}$,<br>rph-1, $\Delta(\text{rhaD-rhaB})568$, hsdR514 | p139 (139,191 bp) | Heidarian et al., 2026 |
| DA81967 | *recC* deletion | F-, $\Delta(\text{araD-araB})567$,<br>$\Delta\text{lacZ4787}(\text{::rrnB-3})$, $\lambda^-$, $\Delta\text{recC747}$,<br>rph-1, $\Delta(\text{rhaD-rhaB})568$, hsdR514 | p139 (139,191 bp) | Heidarian et al., 2026 |
| DA81968 | *recD* deletion | F-, $\Delta(\text{araD-araB})567$,<br>$\Delta\text{lacZ4787}(\text{::rrnB-3})$, $\lambda^-$, $\Delta\text{recD744}$,<br>rph-1, $\Delta(\text{rhaD-rhaB})568$, hsdR514 | p139 (139,191 bp) | Heidarian et al., 2026 |
| DA81797 | KEIO wild-type | Eco wt ME9062 BW25113 rrnB<br>$\Delta\text{lacZ4787}$ hsdR514 $\Delta(\text{araBAD})567$<br>$\Delta(\text{rhaBAD})568$ rph-1" | p178 (178,078 bp) | Heidarian et al., 2026 |
| DA81758 | *recA* deletion | F-, $\Delta(\text{araD-araB})567$,<br>$\Delta\text{lacZ4787}(\text{::rrnB-3})$, $\lambda^-$, $\Delta\text{recA774}$,<br>rph-1, $\Delta(\text{rhaD-rhaB})568$, hsdR514 | p178 (178,078 bp) | Heidarian et al., 2026 |
| DA81759 | *recB* deletion | F-, $\Delta(\text{araD-araB})567$,<br>$\Delta\text{lacZ4787}(\text{::rrnB-3})$, $\lambda^-$, $\Delta\text{recB745}$,<br>rph-1, $\Delta(\text{rhaD-rhaB})568$, hsdR514 | p178 (178,078 bp) | Heidarian et al., 2026 |
| DA81760 | *recC* deletion | F-, $\Delta(\text{araD-araB})567$,<br>$\Delta\text{lacZ4787}(\text{::rrnB-3})$, $\lambda^-$, $\Delta\text{recC747}$,<br>rph-1, $\Delta(\text{rhaD-rhaB})568$, hsdR514 | p178 (178,078 bp) | Heidarian et al., 2026 |
| DA81761 | *recD* deletion | F-, $\Delta(\text{araD-araB})567$,<br>$\Delta\text{lacZ4787}(\text{::rrnB-3})$, $\lambda^-$, $\Delta\text{recD744}$,<br>rph-1, $\Delta(\text{rhaD-rhaB})568$, hsdR514 | p178 (178,078 bp) | Heidarian et al., 2026 |
| DA84878 | KEIO wild-type | Eco wt ME9062 BW25113 rrnB<br>$\Delta\text{lacZ4787}$ hsdR514 $\Delta(\text{araBAD})567$<br>$\Delta(\text{rhaBAD})568$ rph-1" | pminiF (8,803 bp) | This study |
| DA84967 | *recA* deletion | F-, $\Delta(\text{araD-araB})567$,<br>$\Delta\text{lacZ4787}(\text{::rrnB-3})$, $\lambda^-$, $\Delta\text{recA774}$,<br>rph-1, $\Delta(\text{rhaD-rhaB})568$, hsdR514 | pminiF (8,803 bp) | This study |
| DA84971 | *recB* deletion | F-, $\Delta(\text{araD-araB})567$,<br>$\Delta\text{lacZ4787}(\text{::rrnB-3})$, $\lambda^-$, $\Delta\text{recB745}$,<br>rph-1, $\Delta(\text{rhaD-rhaB})568$, hsdR514 | pminiF (8,803 bp) | This study |
| DA84975 | *recC* deletion | F-, $\Delta(\text{araD-araB})567$,<br>$\Delta\text{lacZ4787}(\text{::rrnB-3})$, $\lambda^-$, $\Delta\text{recC747}$,<br>rph-1, $\Delta(\text{rhaD-rhaB})568$, hsdR514 | pminiF (8,803 bp) | This study |
| DA84890 | *recD* deletion | F-, $\Delta(\text{araD-araB})567$,<br>$\Delta\text{lacZ4787}(\text{::rrnB-3})$, $\lambda^-$, $\Delta\text{recD744}$,<br>rph-1, $\Delta(\text{rhaD-rhaB})568$, hsdR514 | pminiF (8,803 bp) | This study |

1. Heyman, G. *et al.* Prevalence, misclassification, and clinical consequences of the heteroresistant phenotype in *Escherichia coli* bloodstream infections in patients in Uppsala, Sweden: a retrospective cohort study. *Lancet Microbe* **6**, 101010 (2025). 2. Heidarian, S., Hjort, K., Nicoloff, H. & Andersson, D. I. Deletions of recombination genes impair tandem amplification and reshape heteroresistance mechanisms in *Escherichia coli*. *mBio* **17**, (2026).

**Table S2.** List of TZP-resistant mutants isolated in this study, with their respective parental strains and mutations. The last column represents the percentage of multimer reads divided by (monomer + multimer) reads only. n = 2 for the parental strains and n = 1 for each mutant tested. ND: not determined

| Strain ID | Genetic background | Mutations | % of multimers |
| --- | --- | --- | --- |
| DA62886 | Clinical Parental | - | 0.23 ± 0.02 |
| DA84058 | DA62886 | Duplication on RNAI-RNAII | 0.07 |
| DA84124 | DA62886 | Duplication on RNAI-RNAII | 0.11 |
| DA84181 | DA62886 | SNP on RNAI-RNAII | 0.22 |
| DA84190 | DA62886 | SNP on RNAI-RNAII | 0.15 |
| DA84249 | DA62886 | Duplication on RNAI-RNAII | 0.08 |
| DA84251 | DA62886 | Duplication on RNAI-RNAII | 0.03 |
| DA84119 | DA62886 | SNP on RNAI-RNAII | ND |
| DA84131 | DA62886 | Duplication on RNAI-RNAII | ND |
| DA84132 | DA62886 | Duplication on RNAI-RNAII | ND |
| DA84139 | DA62886 | 1.Duplication on RNAI-RNAII<br>2.SNP on chromosomal *yhhM* | ND |
| DA84140 | DA62886 | SNP on RNAI-RNAII | ND |
| DA84179 | DA62886 | Duplication on RNAI-RNAII | ND |
| DA84180 | DA62886 | SNP on RNAI-RNAII | ND |
| DA84186 | DA62886 | Duplication on RNAI-RNAII | ND |
| DA84192 | DA62886 | Duplication on RNAI-RNAII | ND |
| DA84197 | DA62886 | Duplication on RNAI-RNAII | ND |
| DA84199 | DA62886 | Duplication on RNAI-RNAII | ND |
| DA84203 | DA62886 | 1.Duplication on RNAI-RNAII<br>2.SNP on chromosomal intergenic region | ND |
| DA84250 | DA62886 | Duplication on RNAI-RNAII | ND |
| DA82535 | MG1655 Parental | - | 0.13 ± 0.18 |
| DA84157 | DA82535 | 1.Frameshift mutation on *recD*<br>2.SNP on chromosomal *iap* gene | 6.14 |
| DA84214 | DA82535 | InDel on RNAI-RNAII | 0.14 |
| DA84222 | DA82535 | 1.Nonsense mutation on *recD*<br>2.SNP on chromosomal *yhjJ* gene | 4.08 |
| DA84234 | DA82535 | Duplication on RNAI-RNAII | 0.00 |
| DA84258 | DA82535 | Duplication on RNAI-RNAII | 0.04 |
| DA84266 | DA82535 | Duplication on RNAI-RNAII | 0.02 |
| DA84154 | DA82535 | InDel on RNAI-RNAII | ND |
| DA84210 | DA82535 | 1.Duplication on RNAI-RNAII<br>2.Tn3 insertion from p12 to chromosomal<br>*ydeA* gene | ND |
| DA84213 | DA82535 | 1.Duplication on RNAI-RNAII<br>2.Tn3 insertion from p12 to chromosomal<br>*ydeA* gene | ND |
| DA84217 | DA82535 | 1.Nonsense mutation on *recD*<br>2.SNP on chromosomal *yhjJ* gene | ND |
| DA84225 | DA82535 | Duplication on RNAI-RNAII | ND |
| DA84228 | DA82535 | Duplication on RNAI-RNAII | ND |
| DA84239 | DA82535 | Duplication on RNAI-RNAII | ND |
| DA84252 | DA82535 | 1.SNP on RNAI-RNAII<br>2.SNP on chromosomal *yhjJ* gene | ND |
| DA84254 | DA82535 | 1.SNP on RNAI-RNAII<br>2.SNP on chromosomal *yhjJ* gene | ND |
| DA84262 | DA82535 | Duplication on RNAI-RNAII | ND |
| DA84269 | DA82535 | Duplication on RNAI-RNAII | ND |
| DA84270 | DA82535 | Duplication on RNAI-RNAII | ND |
| DA84272 | DA82535 | Duplication on RNAI-RNAII | ND |

**Table S3.** Revertant analysis for p12 plasmid loss. Strains DA82535 (parental) and DA84154 (InDel) are shown as controls.

| Strain ID | Mutation type | Colonies tested | % of revertants that have lost the p12 |
| --- | --- | --- | --- |
| DA82535 | parental | 4 | 0% |
| DA84154 | InDel | 6 | 0% |
| DA84157 | *recD* | 18 | 16.67% |
| DA84217 | *recD* | 18 | 22.22% |
| DA84222 | *recD* | 18 | 33% |

**Table S4.**
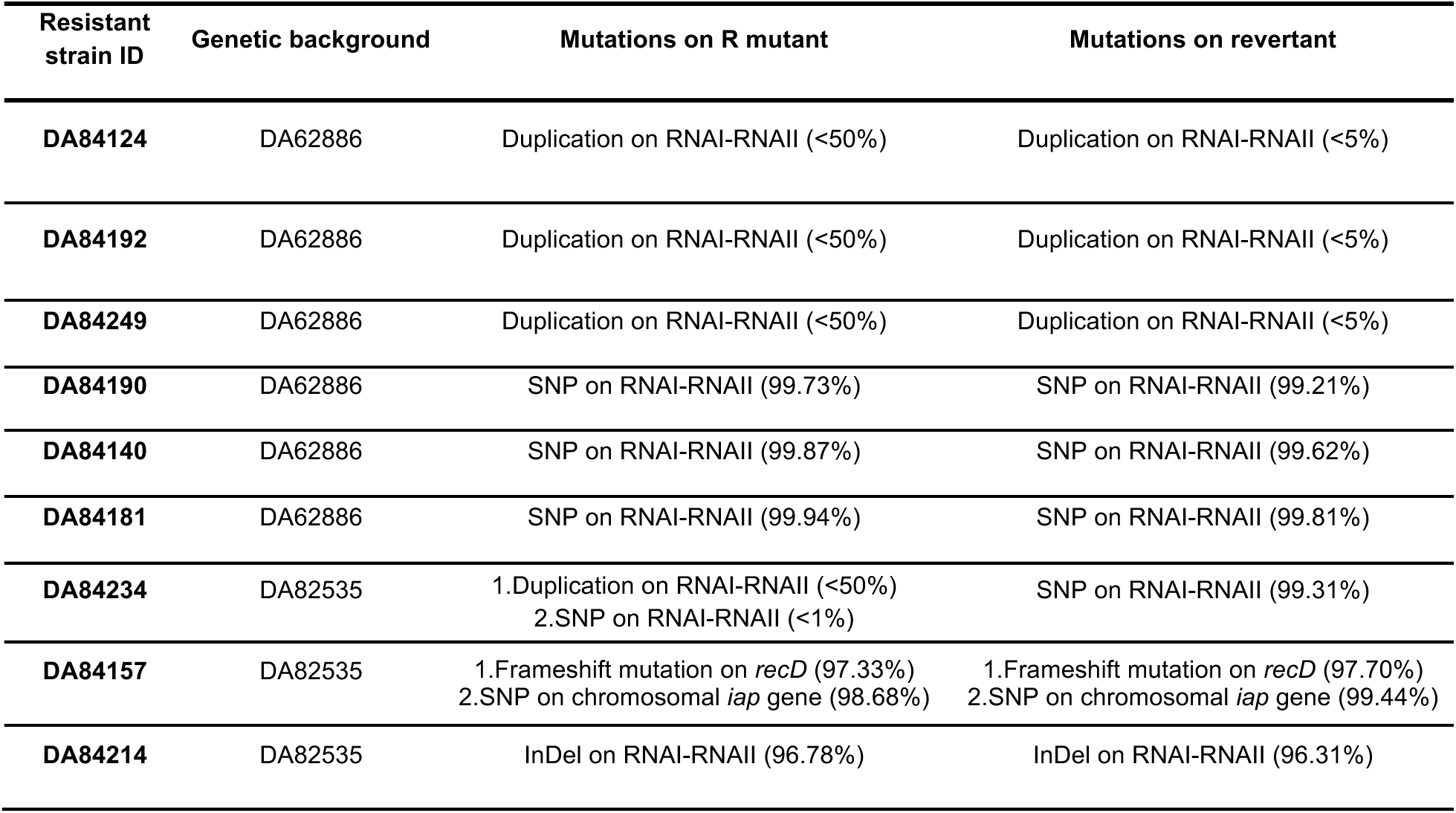
List of mutations of TZP-resistant mutants and a corresponding revertant clone, along with % of frequency of each mutation. For the duplication mutations, the frequency is a rough estimation based on manual inspection (Supplementary Fig. 2).

| Resistant strain ID | Genetic background | Mutations on R mutant | Mutations on revertant |
| --- | --- | --- | --- |
| DA84124 | DA62886 | Duplication on RNAI-RNAII (<50%) | Duplication on RNAI-RNAII (<5%) |
| DA84192 | DA62886 | Duplication on RNAI-RNAII (<50%) | Duplication on RNAI-RNAII (<5%) |
| DA84249 | DA62886 | Duplication on RNAI-RNAII (<50%) | Duplication on RNAI-RNAII (<5%) |
| DA84190 | DA62886 | SNP on RNAI-RNAII (99.73%) | SNP on RNAI-RNAII (99.21%) |
| DA84140 | DA62886 | SNP on RNAI-RNAII (99.87%) | SNP on RNAI-RNAII (99.62%) |
| DA84181 | DA62886 | SNP on RNAI-RNAII (99.94%) | SNP on RNAI-RNAII (99.81%) |
| DA84234 | DA82535 | 1.Duplication on RNAI-RNAII (<50%)<br>2.SNP on RNAI-RNAII (<1%) | SNP on RNAI-RNAII (99.31%) |
| DA84157 | DA82535 | 1.Frameshift mutation on *recD* (97.33%)<br>2.SNP on chromosomal *iap* gene (98.68%) | 1.Frameshift mutation on *recD* (97.70%)<br>2.SNP on chromosomal *iap* gene (99.44%) |
| DA84214 | DA82535 | InDel on RNAI-RNAII (96.78%) | InDel on RNAI-RNAII (96.31%) |

**Table S5.**
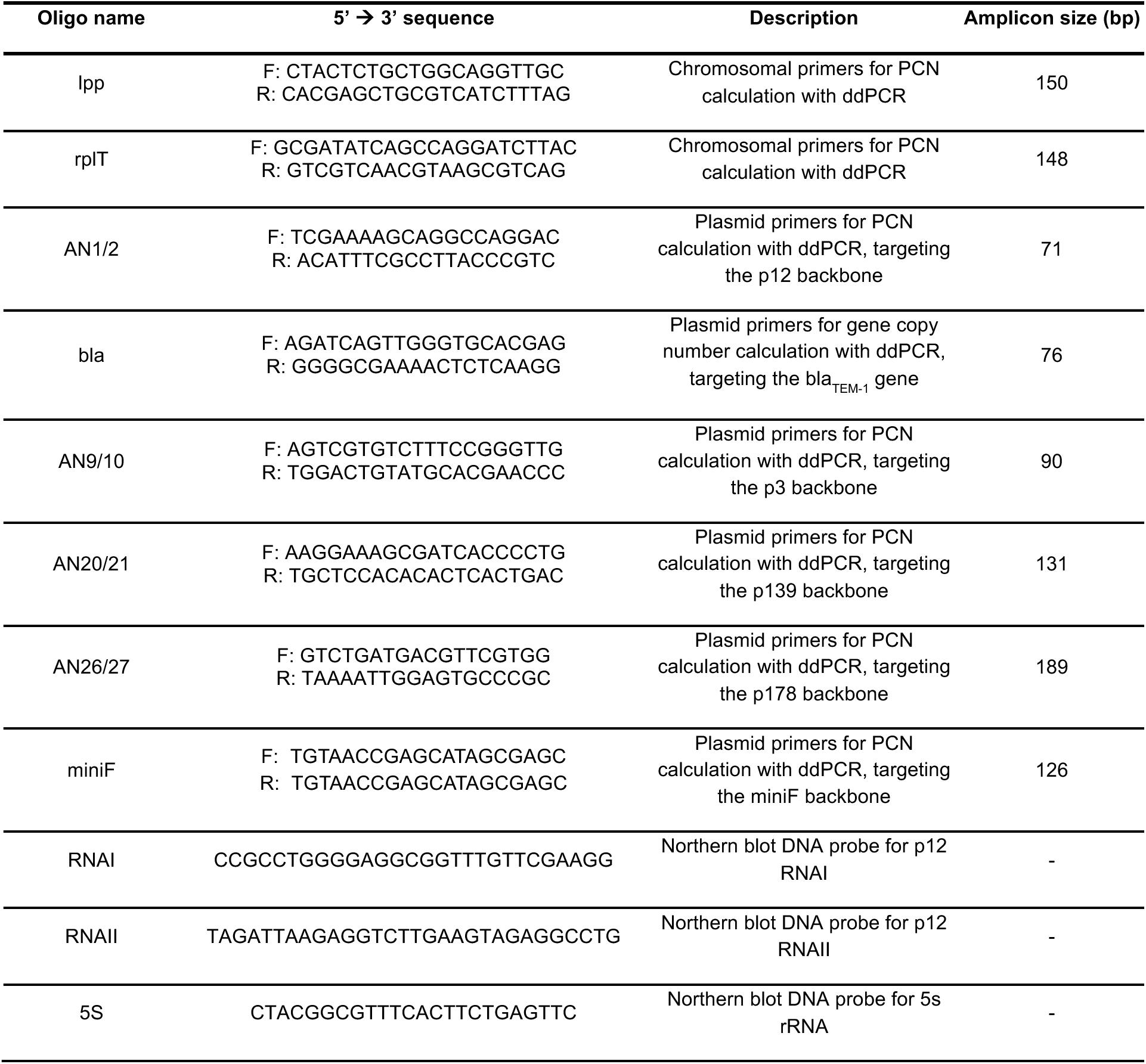
List of oligonucleotides used in this study.

